# A remarkable expansion of secreted cysteine knot proteins reveals novel virulence factors in the human fungal pathogen *Histoplasma*

**DOI:** 10.1101/2025.05.13.653811

**Authors:** Rosa A. Rodriguez, Dinara Azimova, Mark Voorhies, Bevin C. English, Jane Symington, Anita Sil

## Abstract

Identifying fungal secreted factors that influence host infection remains a key challenge in microbial pathogenesis. While secreted effectors, particularly small cysteine-rich proteins, are well characterized in plant fungal pathogens, their counterparts in mammalian pathogens remain elusive. We applied criteria from plant fungal effectors to the mammalian fungal pathogen *Histoplasma*, yielding a set of putative effectors highly enriched for knottins, proteins that adopt a distinctive cysteine knot fold. Using a novel algorithm, we further identified 25 putative knottins in the *Histoplasma* genome, revealing a significant expansion of knottin genes. Knottin domains are found in diverse molecules but play an unknown role in virulence. Functional studies of individual *Histoplasma* knottins demonstrated their critical roles in intracellular survival and host cell lysis during macrophage infection as well as virulence *in vivo*. These findings highlight the importance of knottins in fungal pathogenesis and suggest their broader relevance for discovering conserved mechanisms of host manipulation.

## Introduction

Intracellular pathogens manipulate the environment inside their host and cause death by employing the use of secreted virulence factors^1–3^. Virulence factors, also called effectors, can be molecules, toxins, or proteins used by an organism to attach to or invade host cells, inhibit or evade the host immune system, or promote cell death^4, 5^. Existing paradigms about how effectors function and manipulate the host to create a replicative niche largely come from bacterial pathogens^6–10^. Far less is known about the drivers of pathogenesis in eukaryotic organisms^11^. Identifying shared characteristics of virulence factors can be a powerful approach to expand our understanding of fungal virulence, which is critical given the global increase in fungal infections^12, 13^.

*Histoplasma* is a thermally dimorphic human fungal pathogen and the causative agent of histoplasmosis^14, 15^. In the United States, *Histoplasma* is endemic to the Ohio and Mississippi River Valley regions^16, 17^. In the soil, *Histoplasma* grows as multicellular hyphae and spores^18^. When the soil is disrupted, the hyphal fragments and spores are aerosolized and inhaled by mammals where body temperature of 37°C is sufficient to induce a morphological transition to budding yeast. In the lungs, *Histoplasma* interacts with phagocytic cells like alveolar macrophages where it is readily phagocytosed but not killed^19–21^. *Histoplasma* avoids detection and clearing by the immune system and ultimately induces host cell death, although the exact mechanisms through which this is achieved are still poorly understood^22–25^.

There are a handful of virulence factors that have been shown to be necessary for *Histoplasma* virulence^26–32^, the most well studied of which is Cbp1^33–36^. In previous work we demonstrated that Cbp1 is necessary for lysis of macrophages through the induction of an intracellular signaling cascade called the integrated stress response, or ISR^37^. If the stress remains unresolved, the ISR leads to host cell death via caspase-3/7-mediated apoptosis^38–40^. Cbp1 was observed to localize to the host cytosol^41^, but the mechanisms through which Cbp1 accesses the cytosol or induces the ISR remain unknown. In this work we hope to expand the repertoire of known secreted virulence factors from *Histoplasma*.

As inspiration, we turned to the plant fungal pathogen field, which has identified virulence factors with shared characteristics^42–45^. Many virulence factors are small proteins (< 300 amino acids) that are secreted extracellularly, highly expressed in the pathogenic form of the fungus, and contain enriched cysteine content^11, 46^. The cysteines form disulfide bonds which are thought to confer stability and maintain protein structures in the changing environments the virulence factor may encounter^47–49^. In this work, we aimed to identify novel virulence factors by taking a comprehensive bioinformatic approach and applying some of these criteria to generate a list of candidate virulence factors in *Histoplasma*. We identified predicted proteins that are small, secreted, and contain increased cysteine content, including those that are preferentially expressed by *Histoplasma* yeast cells. This analysis revealed 15 putative effectors, four of which contained homology to a cysteine knot gene family called knottins. The fungal knottins include *AVR9* (*avirulence gene 9*), an effector from the plant fungal pathogen *Cladosporium fulvum.* Avr9 can elicit a plant immune defense response but the mechanisms through which it causes host cell death are still unknown^48, 50–52^. By writing an algorithm to rigorously identify the knottin motif across the *Histoplasma* genome, we uncovered 25 putative knottins and showed that this significant expansion of knottins in *Histoplasma* species is not observed in most other fungi. Furthermore, characterization of four knottins revealed localization to the host cytosol and critical virulence phenotypes in macrophage lysis and intracellular growth. Lastly, we queried two of the knottins in the mouse model of infection and discovered that they are essential for virulence. This work demonstrates a robust methodology to identify effectors that play a key role in *Histoplasma* pathogenesis.

## Results

### Bioinformatic analysis identifies putative *Histoplasma* virulence factors

The fungal effectors that contribute to *Histoplasma* virulence are largely unknown, and only a handful of effectors have been identified^26–32^. We decided to take a bioinformatic approach using existing datasets to uncover novel effectors in *Histoplasma*, basing our analysis on features of previously identified effectors of plant fungal pathogens^11, 42, 46^. We focused on four criteria: small (≤ 250 amino acids), secreted (Phobius prediction)^53, 54^, potential to form disulfide bonds (≥ 4 cysteines), and expression in the pathogenic yeast phase (Fig. 1A). For this latter criterion, we used previously published transcriptomic data of steady-state yeast and hyphae to identify predicted transcripts that are preferentially expressed in the yeast form^55^. Additionally, since Ryp transcription factors are key regulators of yeast-phase growth and directly associate upstream of known effectors like Cbp1^56^, we also determined which putative effectors are direct targets of Ryps. In total, we identified approximately 9580 *Histoplasma* transcripts of which 4323 (45.1%) are predicted to encode proteins that are ≤ 250 amino acids in length, 927 (9.7%) are predicted to encode proteins that are secreted, and 5221 (54.5%) encode proteins that contain at least four cysteines (which we are calling “cysteine-rich”). Of these transcripts, 226 are thought to encode proteins that are small, secreted, and cysteine-rich (Fig. 1B). Approximately 683 (7.1%) predicted genes are Ryp targets and 1271 (13.3%) display yeast-enriched expression. By examining the intersection of these criteria, we identified 15 putative effectors (Fig. 1C, Fig. S1). Importantly, this list contains three previously identified virulence factors (Cbp1^26, 36^, Sod3^27^, and Yps3^28, 41^), providing proof of principle that our bioinformatic approach can accurately identify *Histoplasma* effectors. Moreover, two hits (the paralogs YPS21^57^ and GH17/CFP4^58–60^) encode secreted yeast-enriched factors with undefined roles in virulence. Seven other hits are *Histoplasma* proteins of unknown function with no predicted domains. Finally, the remaining three were identified in our previous work as containing homology to the Fungi1 knottin family, a protein family with a conserved cysteine knot motif^50, 51, 55^ that has not previously been explored for roles in pathogenesis in *Histoplasma*.

**Figure 1.**
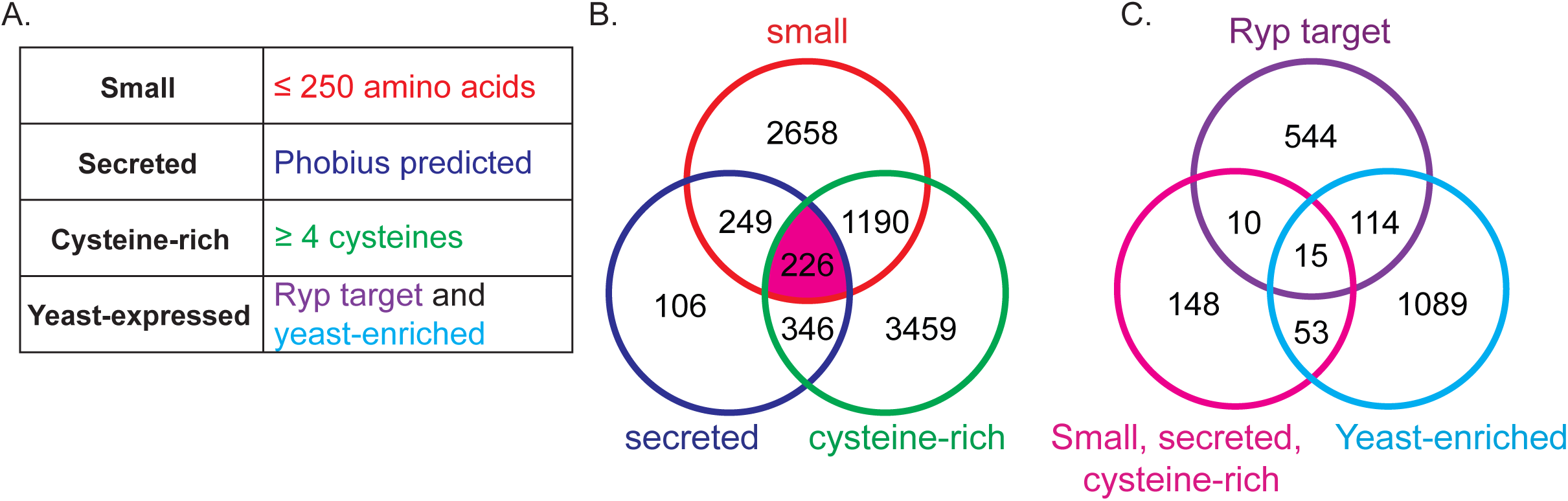
Criteria for identifying novel virulence factors in *Histoplasma*. (A) Candidate proteins were required to be small (≤ 250 amino acid), secreted (signal sequence predicted by Phobius), cysteine-rich (≥ 4 cysteines), and yeast-expressed (contain at least one upstream Ryp-binding event and enriched in the yeast form as measured by RNA-seq). (B) Venn diagram identifying 226 candidate proteins that were small, secreted, and cysteine-rich. *Histoplasma* contains 9,580 protein coding transcripts of which 4,323 are predicted to encode proteins less than 250 amino acids in length, 927 are predicted to be secreted, and 5,221 contain at least four cysteines. (C) Venn diagram identifying 15 protein-coding genes that were Ryp targets and preferentially expressed in the yeast form. In total, 683 transcripts were direct targets of at least one Ryp (Ryp1-4) transcription factor, and 1,271 were preferentially expressed in the yeast form of *Histoplasma*.

### Knottins are greatly expanded in *Histoplasma* species and not in other closely related fungi

We were intrigued by the presence of three knottin proteins in the set of putative *Histoplasma* effectors. Knottin domains are characterized by the presence of at least six cysteines that form disulfide bonds in an interwoven configuration, generating a knot-like structure that confers resistance to heat and proteolysis^50, 51, 61^. Knottins are found in plants, animals, insects, and fungi and proteins containing this domain have extremely diverse functions including but not limited to protease inhibitors^62^, insecticides^63, 64^, a human neuropeptide^65^, and ion channel inhibitors^66^. In fungi, knottin proteins fall into two distinct groups, with the secreted proteins belonging to the Fungi1 family^45, 50^ including Avr9. Hence, we were interested in the possibility that knottin proteins might play important roles in *Histoplasma* biology. We first mined the *Histoplasma* genome to determine whether other knottin proteins were present. During our previous annotation of the *Histoplasma* transcriptome, we identified 12 putative knottins in the *Histoplasma* G217B genome that contained the conserved gene structure from plant and fungal knottins^55^. To find more knottins, we focused on the gene structure of fungal knottins, where it has been observed that knottins are comprised of two exons, with the six-cysteine knottin motif in the second exon^55, 67^. Additionally, for the *Histoplasma* G217B strain, the knottin motif has highly conserved spacing between the cysteines (Fig. 2A).

**Figure 2.**
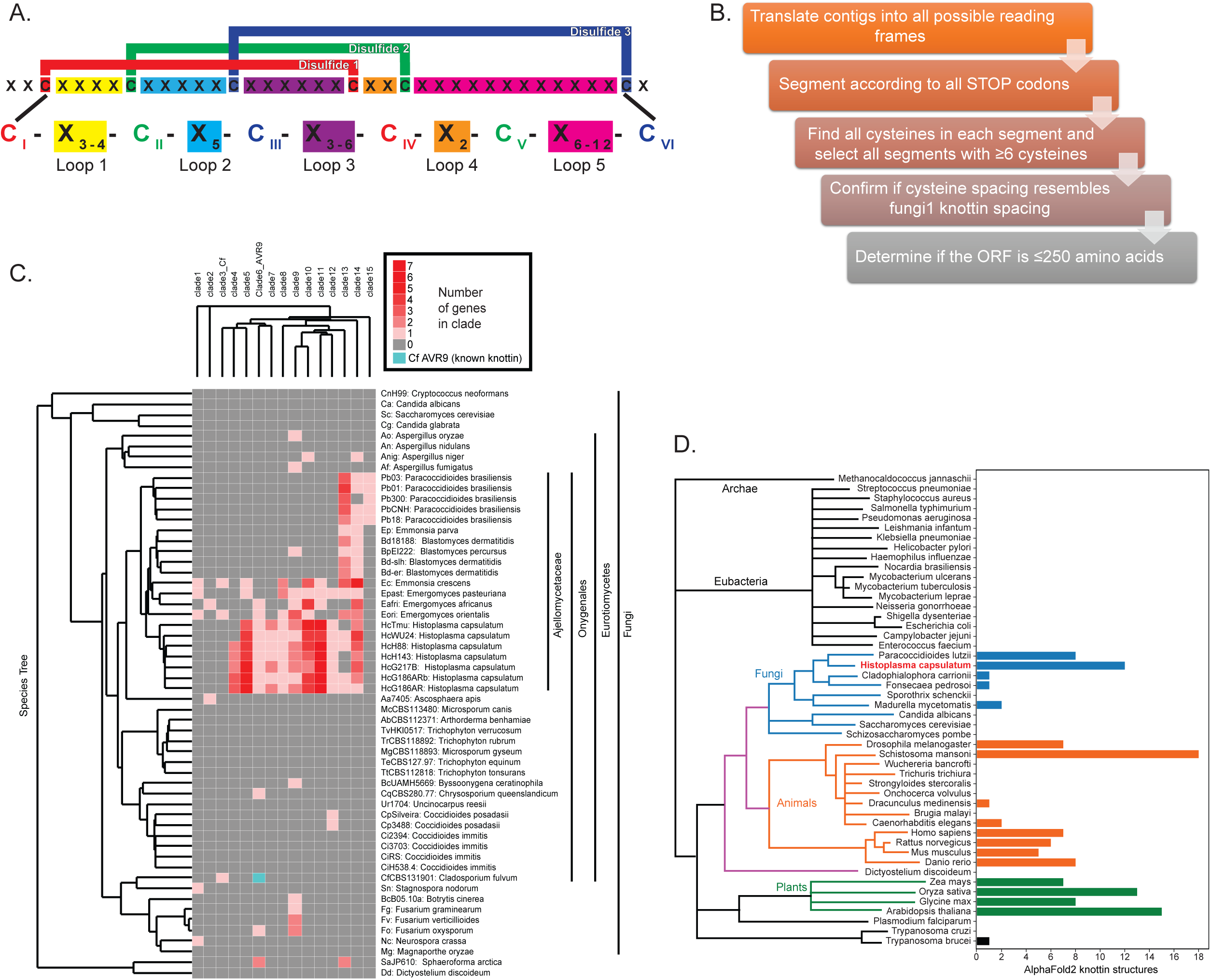
Knottin proteins are expanded in *Histoplasma* and not in other closely related fungi. (A) Schematic of the knottin motif in *Histoplasma* G217B strain. The canonical fungal knottin motif consists of at least 6 cysteines that form three disulfide bonds. The red line denotes the first disulfide bond formed between cysteine 1 and 4, the green line denotes the second disulfide bond between cysteine 2 and 5 and the blue line denotes the third disulfide bond between cysteine 3 and 6. Loop 1 (yellow) is 3-4 amino acids in length, loop 2 (light blue) is 8 amino acids long, loop 3 (purple) is 3-6 amino acids long, loop 4 (orange) is 2 amino acids long and loop 5 (pink) is the most variable in length, ranging from 6-12 amino acids. (B) Diagram of the steps in the KNOTTIN_FINDER algorithm. (C) Phylogenetic analysis of knottins in 56 representative fungal genomes (rows). Knottins cluster into 15 distinct gene clades (columns). The intensity of the red indicates the number of knottin genes present in each clade per genome. Teal square denotes *AVR9* from *Cladosporium fulvum*. Branch lengths are not to scale. (D) Expansion of the knottin motif across 16 model organism proteomes and 32 proteomes from organisms important for global health using AlphaFold2 predictions and KNOTTIN_TOPOLOGY_SCAN algorithm. No knottin containing proteins were found in bacteria.

Based on the observed exon structure, which implies that all the knottin cysteine codons will lie in a single open reading frame (ORF), we designed a novel algorithm (KNOTTIN_FINDER) to detect knottins across the *Histoplasma* genome by searching for the conserved cysteine spacing pattern^55^ (Fig. 2A). Any ORFs ≥ 750 nucleotides were discarded so as to exclude large proteins that may contain many cysteines. Additionally, we removed any mature CDS that was longer than 750 nucleotides (Data S1, Fig. 2B). As proof of principle, we ran this algorithm against the *Cladosporium fulvum* genome, which contains *AVR9* (avirulence gene 9), the founding member of the Fungi1 knottin family^50, 51^. KNOTTIN_FINDER identified both *AVR9* and a second hit annotated as a “putative effector” in an unpublished study (genbank accession AQA29249)^68^.

Scanning the full *Histoplasma* G217B genome with the KNOTTIN_FINDER algorithm identified a total of 25 putative knottin proteins including the 12 knottins we originally identified^55^. To infer whether the predicted coding sequence was actually translated, we plotted these hits against existing ribosome footprint data^55^, revealing signal in the yeast phase that displayed the expected single intron 5’ to the six cysteine knottin motif (Fig. S2A). In at least one case, the knottin predicted by KNOTTIN_FINDER was absent from both the predicted gene set from the original G217B annotation^69, 70^ as well as the RNA-seq-based transcriptome assembly^55^, highlighting both the difficulty of detecting knottins by conventional gene prediction methods and the sensitivity of our method (Fig. S2B). Ribosome profiling data and RNA-seq transcriptome data are shown for all 25 putative knottins (Fig. S1, S3). Some knottins were not identified in our initial analysis (Fig. 1A) because we lack data for their regulation in the yeast phase or they do not meet our other criteria (Fig. S3, S4). We observed that the knottins are randomly distributed across all chromosomes of the *Histoplasma* G217B genome^71^ and show no preference for any one genomic region (Fig. S5). Additionally, we could bin knottins into distinct gene clades on the basis of their primary sequence (Fig. S6). When the genomes of five additional *Histoplasma* genome assemblies (G186AR, H88, H143, WU24, and Tmu) were queried with KNOTTIN_FINDER, 22 to 27 knottins were identified per genome. Therefore, the large number of knottin proteins is consistent across the *Histoplasma* strains, and the larger *Histoplasma* G217B and H88 genome sequences (∼37-40Mb vs. ∼33Mb for the other strains) is not correlated with a larger number of predicted knottins. Taken together with the identification of *AVR9* in *C. fulvum*, these results suggest that KNOTTIN_FINDER is both sensitive and specific.

Since knottins are ubiquitous throughout the tree of life, we hypothesized that the knottin motif would be present in many protein-coding genes across the fungal kingdom, much like our observation in *Histoplasma*. To our surprise, when we ran the KNOTTIN_FINDER algorithm against 623 fungal genomes from the ENSEMBL fungal genomes collection, we discovered only a few knottins. KNOTTIN_FINDER identified ∼0- 5 knottins in most fungal genomes, including hits in the basal phyla *Mucormycota* and *Chytridiomycota*, suggesting an ancient origin of the Fungi1 family. The *Histoplasma*- containing family *Ajellomycetaceae* was striking in the significant expansion of knottin hits relative to the other genomes in this search.

To further investigate the expansion in the *Ajellomycetaceae*, we prepared a select set of 56 genomes. This included the 41 genomes from our previous protein-level HMM (hidden Markov model) search^55^, 10 additional genomes from *Ajellomycetaceae*, 3 additional genomes from *Onygenaceae* (a sister family to *Ajellomycetaceae*, containing the thermally dimorphic fungal pathogen *Coccidioides*); *Cladosporium fulvum* and *Ascosphaera apis* (causative agent of the bee disease chalk brood), which is a basal member of *Onygenales* (the order containing *Ajellomycetaceae* and *Onygenaceae*) (Table 1). Many of these genomes were newly available and not part of the ENSEMBL fungal genomes from the previous search. Additionally, the GenBank records for many of these additional genomes lack protein annotations, presenting difficulties for the protein HMM-based approach we previously employed^55^, but not to our current nucleotide-based approach. When run against these 56 genomes, the KNOTTIN_FINDER algorithm recapitulated our previous result, showing the expansion of the knottins specifically in *Histoplasma* (Fig. 2C). *Paracoccidioides*, another intracellular thermally dimorphic fungal pathogen and in the oldest diverging branch of *Ajellomycetaceae*^72^, has 4-6 knottins per genome, suggesting an expansion at the root of the *Ajellomycetaceae*, perhaps coincident with loss of keratinase activity^73, 74^ and gain of an intracellular niche. The genus *Blastomyces* (including *Emmonsia parva*), has 2-3 knottins per genome. *Emergomyces,* a newly identified genus containing thermally dimorphic species with either yeast or adiaspore morphologies^75, 76^, and sister to *Histoplasma* and *Blastomyces*, has 10-17 knottins per genome. Taken together with the *Emergomyces* and *Histoplasma* result, this suggests expansion of knottins in the *Histoplasma* lineage following divergence from *Paracoccidioides*, and the specific loss of knottins in the *Blastomyces* lineage may be coincident with the preference for an extracellular niche^72^. Outside of the *Ajellomycetacea*, the *Onygenales* have 0-2 knottins, and outside of the *Onygenales*, few fungal knottin genes were identified (Fig. 2C). The expansion of knottins in the *Histoplasma* genome implies they may play a key role in *Histoplasma* biology.

**Table 1.**
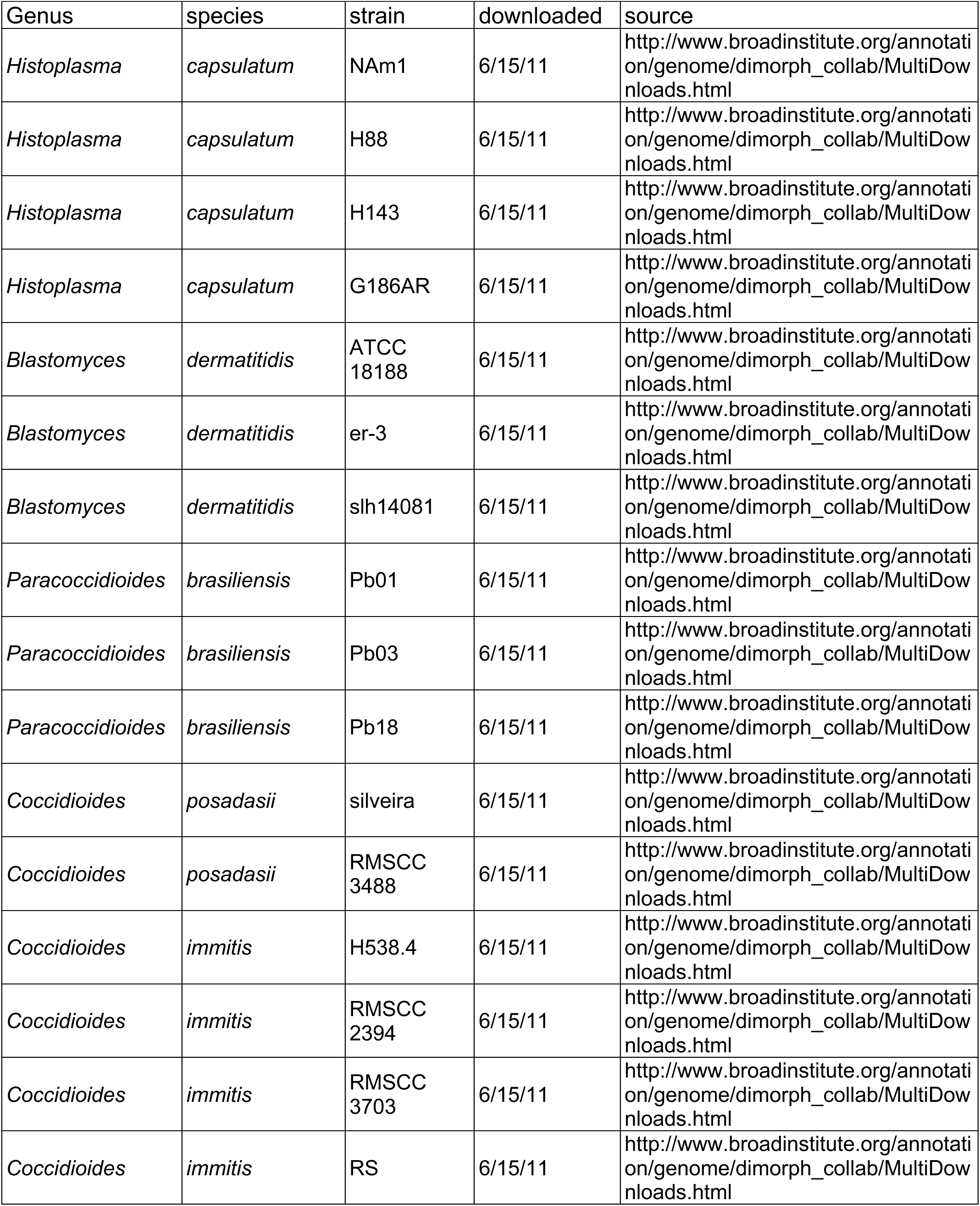
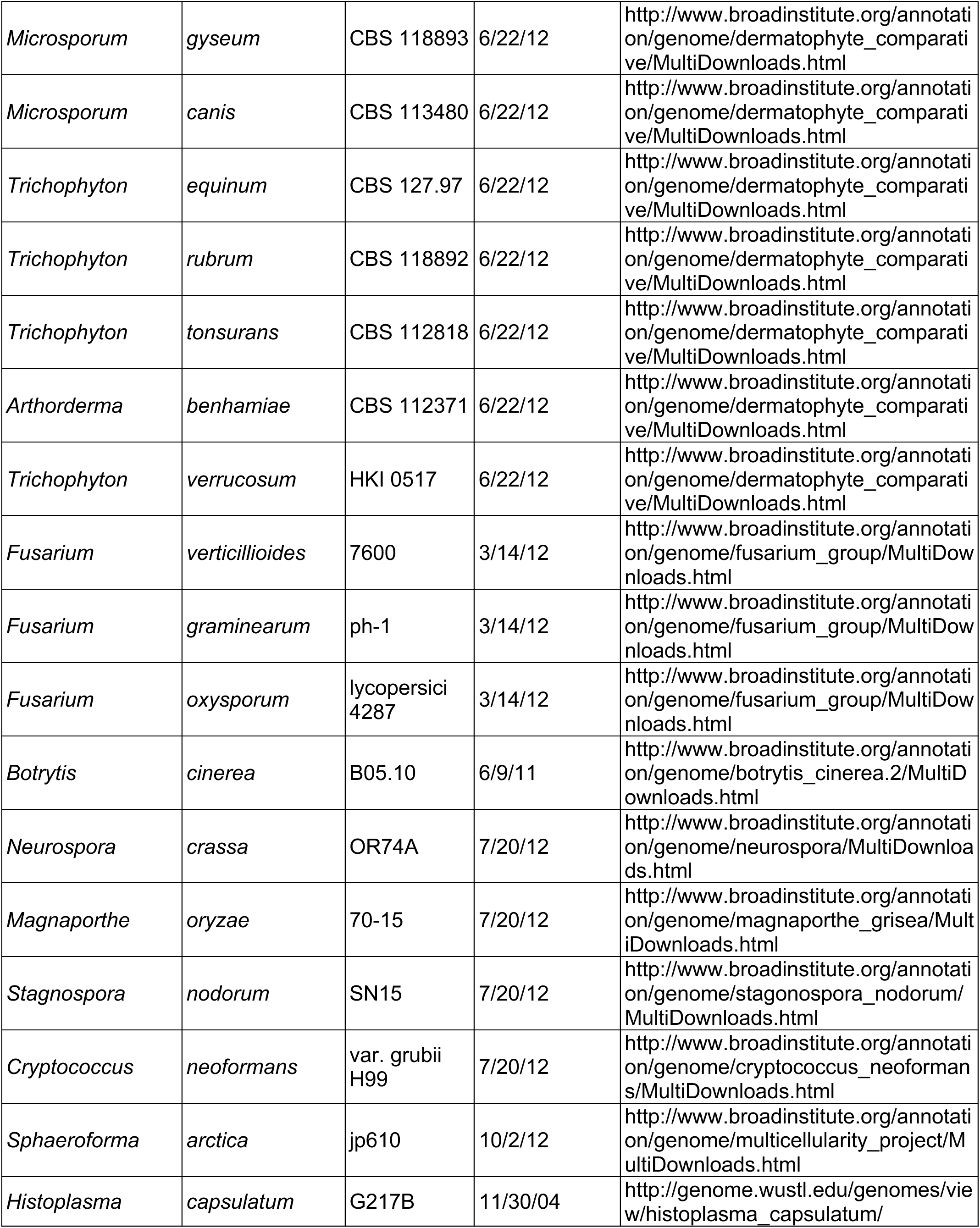
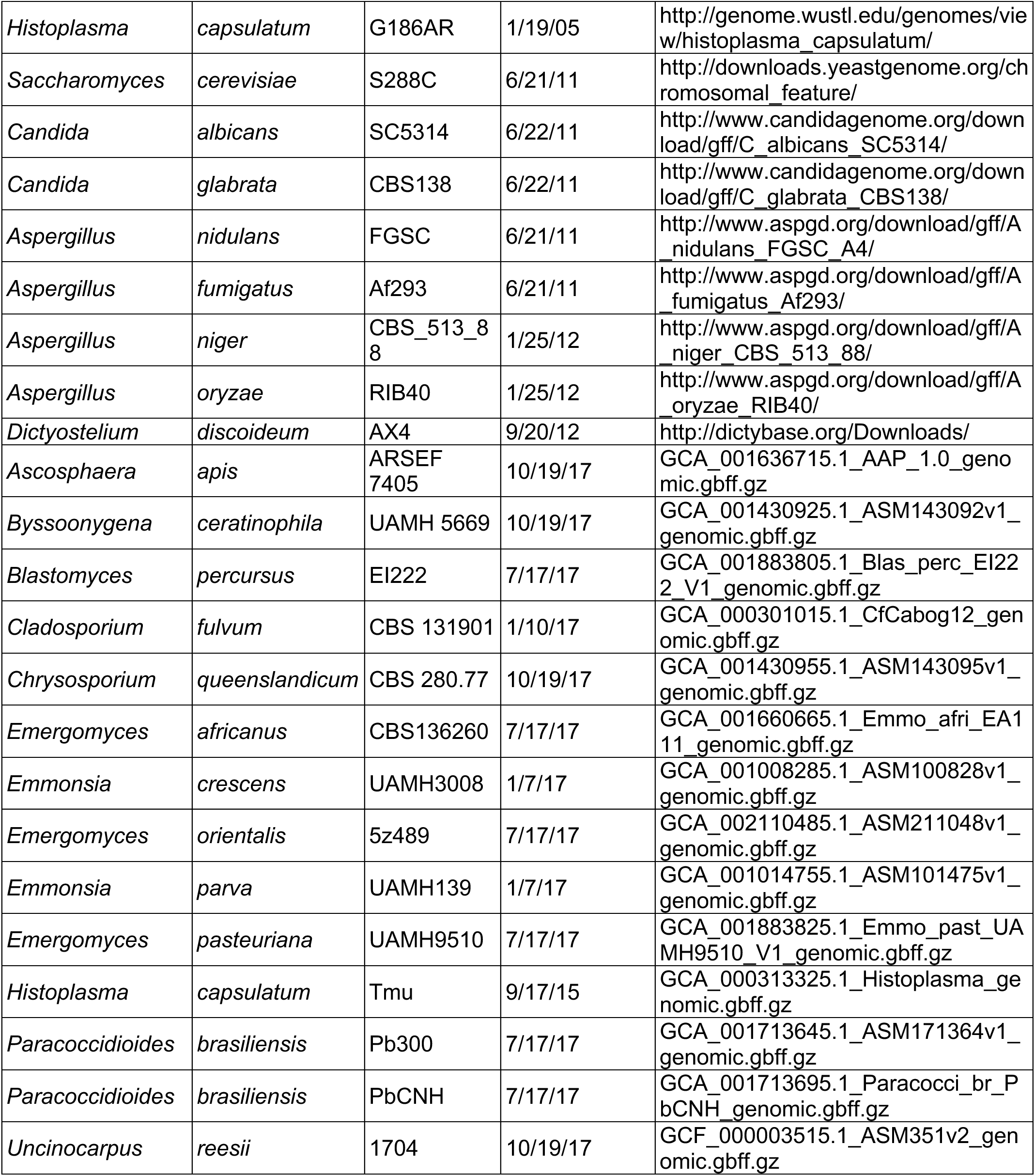
Genomes used to generate phylogenetic tree and identify knottins across different fungal species in Figure 2C.

We decided to employ AlphaFold2^77^ to predict protein structures of this expanded knottin family. For 17 of 25 sequences found by KNOTTIN_FINDER, AlphaFold2 successfully predicted a knottin fold (Fig. S7A). Even though AlphaFold2 did not predict plausible alternative structures for the remaining 8 candidates, given their sequence similarity to the Fungi1 family, we hypothesized that these sequences also adopt a knottin fold. To predict structures for all 25 knottin sequences in a consistent manner, we used Modeller^78^ to generate a homology model for each sequence, using all 17 AlphaFold2 predictions as templates (Fig. S7A). Using these predicted protein structures, we employed APBS^79^ to calculate electrostatic surface potentials for all 25 knottin proteins (Fig. S7B). Of note, surface charges revealed a diversity of charge patterns among the *Histoplasma* knottins. Interestingly, some knottins within the same clade shared certain similarities. For example, clade 14 contains knottins 013, 015 and 020, which are all predicted to have positive surface charges on one face. Analogously, clade 11 knottins are predicted to have negative surface charges on one face. Notably, the predicted knottin structures are diverse and do not reveal specific functional insights.

The successful AlphaFold2 prediction of the knottin fold for 17 out of 25 KNOTTIN_FINDER hits was striking, particularly given that the rarity and high sequence divergence of knottins precludes the deep sequence alignments needed for the covariance matrix used by 80% of AlphaFold2’s neural network. We therefore wondered whether the knottin fold is a ubiquitous false positive prediction generated by AlphaFold2 when presented with a small cysteine-rich protein with few homologs. Alternatively, it may be that the knottin fold is particularly well suited for *de novo* prediction by the physical and geometric layers in the final stage of AlphaFold2’s neural network. To distinguish these cases, we wrote a program (KNOTTIN_TOPOLOGY_SCAN, Data S2, Fig. S8) to scan a set of AlphaFold2 predictions for 48 proteomes (all proteins were run as monomers) generated by UniProt (Table 2)^77^. AlphaFold2 predicted that *Histoplasma* has the highest number of predicted knottins out of the sampled fungal proteomes and failed to predict knottins in *Saccharomyces cerevisiae* or *Candida albicans* (Fig. 2D). Interestingly, no knottin proteins were predicted in any of the bacterial proteomes tested, consistent with bacterial literature lacking reports of knottin proteins^67, 80^. We interpret these findings to mean that the knottin fold is not a common false positive prediction generated by AlphaFold2.

**Table 2.**
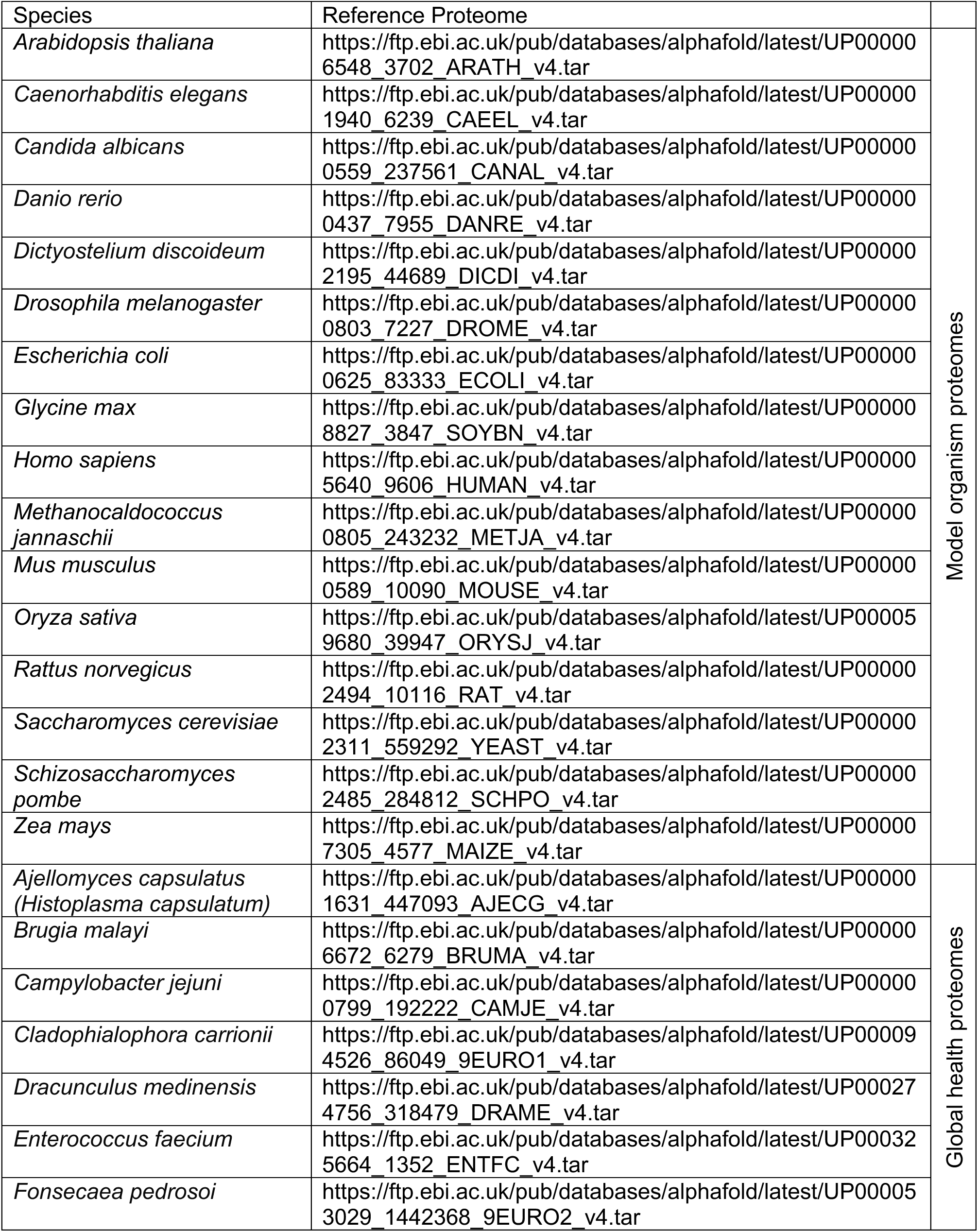
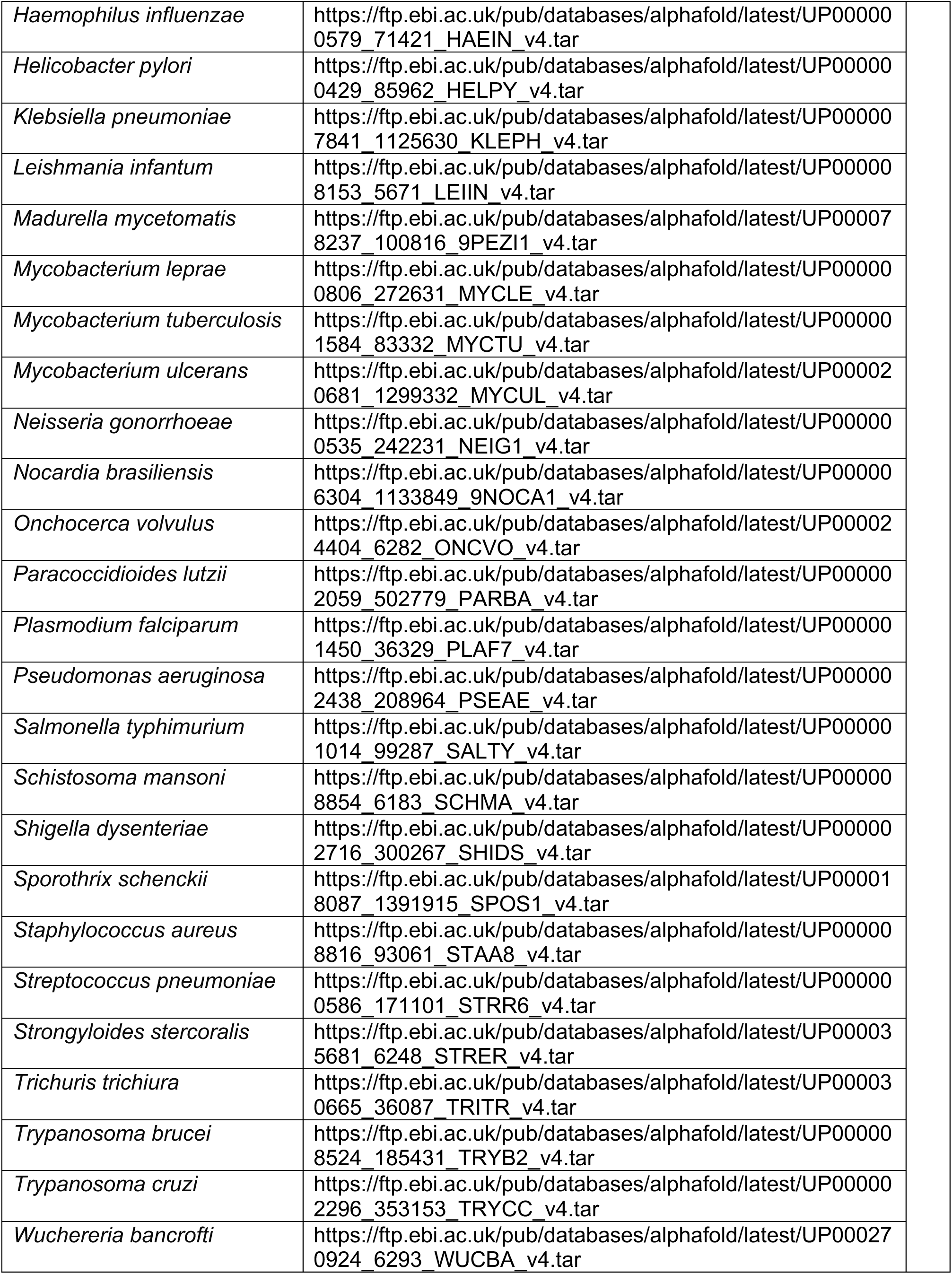
Reference proteomes from model organism and organism implicated as important for global health used in Figure 2D.

### Knot1-4 are required for *Histoplasma* virulence in macrophages

With the expansion of knottins and their expression in the pathogenic yeast form, we hypothesized they may play a role in virulence. Four knottins representing different *Histoplasma* G217B knottin clades were chosen and named *KNOT1* (*KNOTTIN_009*, clade 12)*, KNOT2* (*KNOTTIN_006*, clade 9)*, KNOT3* (*KNOTTIN_023*, clade 11), and *KNOT4* (*KNOTTIN_025*, clade 6) (Fig. S6). Within the *Histoplasma* G217B genome, only one copy of *KNOT1* and *KNOT4* was identified, while *KNOT2* had one paralog (*KNOTTIN_012*) and *KNOT3* had six paralogs (*KNOTTIN_001*, *KNOTTIN_008*, *KNOTTIN_010*, *KNOTTIN_016*, *KNOTTIN_018*, and *KNOTTIN_021*) (Fig. S9).

To directly explore the role of *KNOT1, KNOT2, KNOT3* and *KNOT4* in virulence, we generated deletion mutants using CRISPR-Cas9^81, 82^ and validated gene disruption via PCR, qRT-PCR and genome sequencing (Fig. S10). Complementation strains were generated by expressing the relevant knottin gene under the control of its own promoter and screened for restoration of wild-type expression levels (Fig. 3, Fig. S10D). Although *KNOT1*, *KNOT2*, and *KNOT4* were dispensable for *in vitro* growth of *Histoplasma* yeast cells in liquid cultures, the *knot3Δ* mutant had a slight growth defect (Fig. S11).

**Figure 3.**
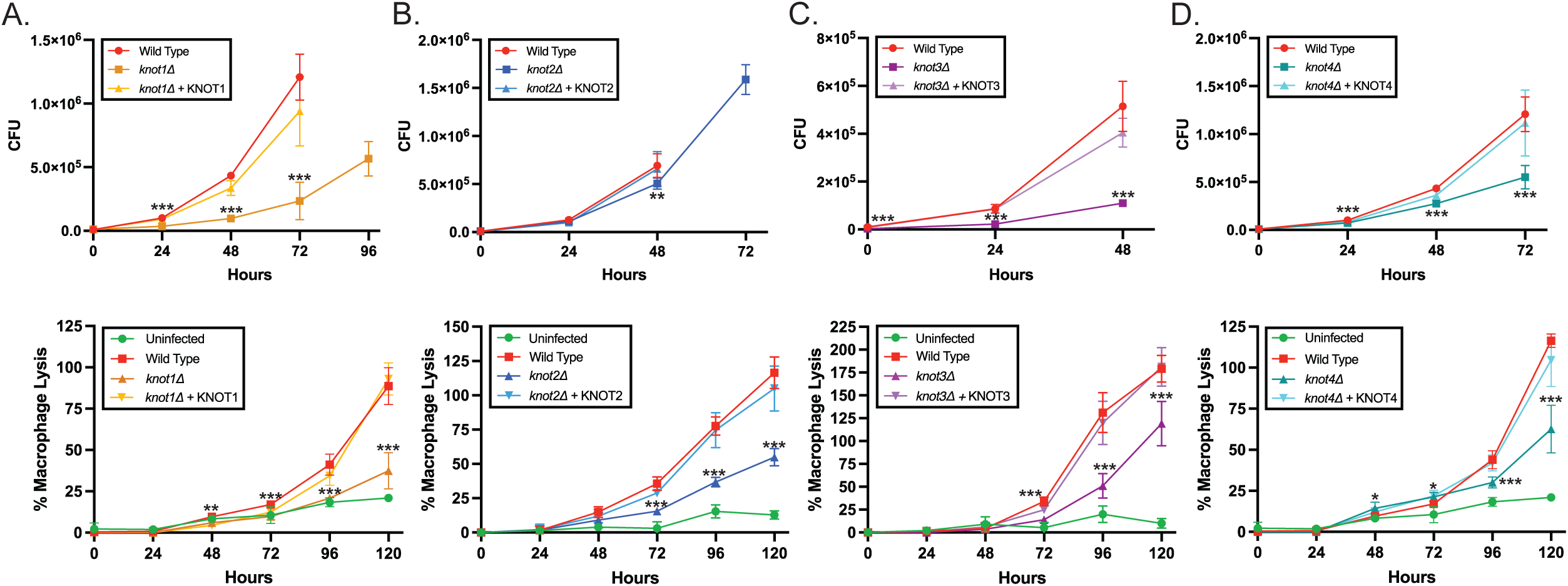
*KNOT1-4* are required for manipulation of macrophages by *Histoplasma.* BMDMs were infected with control or mutant strains at an MOI of 0.5 yeast to 1 macrophage. Lactate dehydrogenase (LDH) assay was used to measure % BMDM lysis over 120 hours. All time points are relative to time point zero of uninfected cells lysed with 1% Triton-X in DMEM (total LDH). Colony forming units (CFU) time points were counted until onset of macrophage lysis to prevent counting extracellular yeast. For CFU and LDH analysis, asterisks indicate p-value < 0.05 relative to wild-type according to t-test. (A) *KNOT1* is required for intracellular growth (top) and macrophage lysis (bottom). (B) *KNOT2* is dispensable for intracellular growth (top) but is required for macrophage lysis (bottom). (C) *KNOT3* is required for intracellular growth (top) and macrophage lysis (bottom). (D) *KNOT4* is required for intracellular growth (top) and macrophage lysis (bottom).

An essential aspect of *Histoplasma* yeast’s pathogenicity is the ability to colonize and lyse macrophages^19–22, 25^, thus, we first assessed whether *KNOT1, KNOT2, KNOT3* and *KNOT4* were required for intracellular growth within bone-marrow derived macrophages (BMDMs). We infected BMDMs at a multiplicity of infection (MOI) of 0.5 yeast to 1 macrophage. Infections were performed with wild-type *Histoplasma*, *knot1Δ, knot2Δ, knot3Δ or knot4Δ* deletion mutants and their respective complementation strains. We quantified intracellular fungal burden by enumerating colony-forming units (CFUs) of *Histoplasma* yeast replicating in macrophages until we observed onset of macrophage lysis and release of intracellular yeast. We discovered that *KNOT1*, *KNOT3* and *KNOT4* were required for optimal intracellular growth whereas *KNOT2* was dispensable (Fig. 3A-D, top panels). The requirement of *KNOT3* for intracellular growth within macrophages may be secondary to a general role in yeast-phase growth, as this mutant displayed a mild growth defect in liquid culture (Fig. S11).

Next, we explored the role of *KNOT1, KNOT2, KNOT3* and *KNOT4* in lysis of host cells. BMDMs were either mock-infected or infected at a MOI of 0.5 with wild-type *Histoplasma*, deletion mutants or the complemented strains. Lactate dehydrogenase (LDH) assays were performed to quantify release of the cytosolic enzyme LDH in supernatants from infected macrophages over the course of the infection. *KNOT1, KNOT2, KNOT3* and *KNOT4* were each required for optimal host cell lysis (Fig. 3A-D, bottom panels). In the case of *KNOT1*, *3*, and *4*, the lysis defect could be secondary to the intracellular growth defect. Since loss of *KNOT2* does not affect intracellular growth *KNOT2* may contribute to host-cell death through a distinct mechanism.

### Knot1-4 access the host cytosol but are not required for ISR activation

To further interrogate the role of knottins in lysis of macrophages by *Histoplasma*, we used subcellular fractionation of infected macrophages to determine the localization of knottins during infection. These studies were modeled on our previous analysis of the *Histoplasma* virulence factor Cbp1, which accesses the host cytosol during infection^41^ and triggers host cell lysis through the induction of the integrated stress response (ISR)^37^. To detect the knottin proteins during infection, we tagged the C-termini of Knot1, Knot2, Knot3, or Knot4 with 3X-FLAG, as knottin proteins are known to be synthesized with an N-terminal propeptide that is cleaved to yield the mature knottin^83, 84^. All four tagged knottins ran at their predicted sizes, ranging from 8-12 kDa. Species corresponding in size to the mature knottin protein (8-12 kDa) as well as the full-length proteins containing both the propeptide and the knottin domain (15-25 kDa) were detected in yeast cell pellets, but the higher molecular weight species, presumably containing the propeptide, were not observed in culture supernatants (Fig. 4A). These data suggest that the cleavage of the propeptide happens before or concomitant with knottin secretion from yeast cells.

**Figure 4.**
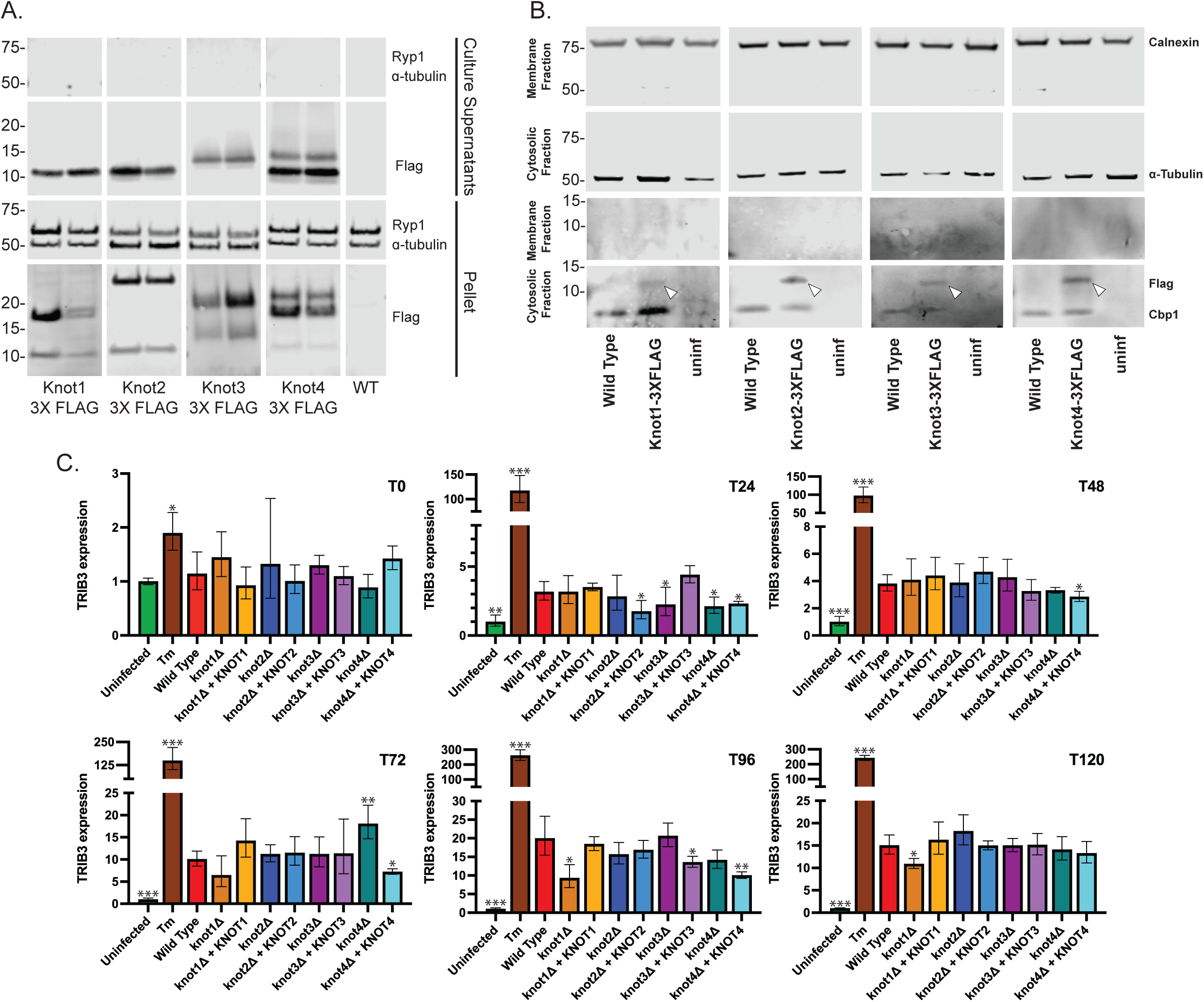
Knottins are secreted by *Histoplasma* and access the macrophage cytosol, but function independently of ISR induction. (A) *Histoplasma* yeast cultures expressing FLAG-tagged knottins were grown and fractionated into culture supernatants and cell pellets. Two knottin species were detected in cell pellets and were inferred to be the full-length protein containing the propeptide and knottin (higher molecular weight band) and a species where the propeptide has been cleaved (lower molecular weight band). In culture supernatants, only the smaller species was detected. Two independent *Histoplasma* colonies expressing Knot1-4 3X-FLAG are shown along with wild-type untagged *Histoplasma*. Ryp1 (*Histoplasma* transcription factor custom antibody)^56^ and ɑ-tubulin antibodies were used as loading controls and FLAG antibody was used to detect FLAG-tagged knottins. (B) BMDMs were mock-infected (uninf) or infected with either the wild-type strain or the FLAG-tagged knottin strains. Macrophage lysates were subjected to subcellular fractionation to separate the cytosolic and membrane fractions followed by SDS-PAGE and Western blot analysis. ɑ-tubulin antibody was used as a marker for the cytosolic fraction and calnexin (endoplasmic reticulum membrane-anchored protein) antibody was used as a marker for the membrane fraction. Cbp1 custom antibody^41^ was used as a positive control. FLAG antibody was used to detect tagged knottins. White triangles denote FLAG-tagged knottins detected in the host cytosol. (C) BMDMs were infected at MOI of 0.5 and RNA was extracted at given time points (0, 24, 48, 72, 96 and 120 hours post infection). qRT-PCR of mouse *TRIB3* expression relative to housekeeping gene mouse *HPRT* is shown. Tunicamycin (Tm, 2.5 μg/mL) was used a positive control for ISR induction. Asterisks indicate p-value < 0.05 relative to wild-type infected BMDMs according to t-test.

Next, we used subcellular fractionation of infected macrophages to determine whether these knottin proteins access the host cytosol as observed previously for Cbp1^41^. Remarkably, Knot1, Knot2, Knot3, and Knot4 were all detected in the host cytosol from infected cells (Fig. 4B). We did not observe any FLAG signal in the membrane fraction, indicating that we could not detect knottin protein associated with the *Histoplasma*-containing phagosome and suggesting that the majority of these knottin proteins are present in the host cytosol.

We next wondered whether knottins were inducing cell death through the macrophage ISR, as observed previously for Cbp1^37^. In brief, the ISR is an intracellular signaling cascade that is activated after exposure to a variety of stresses, including endoplasmic reticulum (ER) stress and amino acid starvation. ISR induction either facilitates return to homeostasis or leads to apoptosis if the stress remains unresolved^38–40^. We previously determined that *Histoplasma* activates the ISR in infected macrophages, as indicated by an increase in eIF2α phosphorylation as well as induction of the transcription factor *CHOP* and the pseudokinase Tribbles 3 (*TRIB3*)^37^.

To assess whether Knot1, Knot2, Knot3, or Knot4 are required to induce the ISR, we measured the expression levels of *TRIB3* during infection as a proxy for ISR activation. We infected BMDMs with each knottin mutant and its corresponding complementation strain. The ability of each knottin mutant to induce *TRIB3* was indistinguishable from wild-type *Histoplasma* yeast (Fig. 4C), indicating that ISR induction was not dependent on each of these four knottins.

### *KNOT2* and *KNOT4* are required for *Histoplasma* virulence *in vivo*

Since *KNOT1-4* were required for colonization and/or lysis of macrophages (Fig. 3), we next wanted to use the murine model of *Histoplasma* infection^26, 37, 41^ to determine their role in pathogenesis. Of the four knottins queried here, we chose to further investigate *KNOT2* because it has a role in macrophage lysis but is dispensable for intracellular growth, which was unique among the four knottins we queried. Additionally, we examined the role of *KNOT4* during *in vivo* infection since *KNOT4* is the ortholog of *AVR9* in *C. fulvum* which has been shown to play a role in plant pathogenesis. C57BL/6J mice were infected intranasally with wild-type, *knot2Δ* or *knot4Δ* mutants, or their respective complementation strains with a lethal dose of *Histoplasma* (1 x 10^6^ yeast). Although mice infected with *knot2Δ* or *knot4Δ* mutants suffered a modest weight loss, they ultimately recovered (Fig. S12), leading us to conclude that both *KNOT2* and *KNOT4* are required for virulence (Fig. 5A and B).

**Figure 5.**
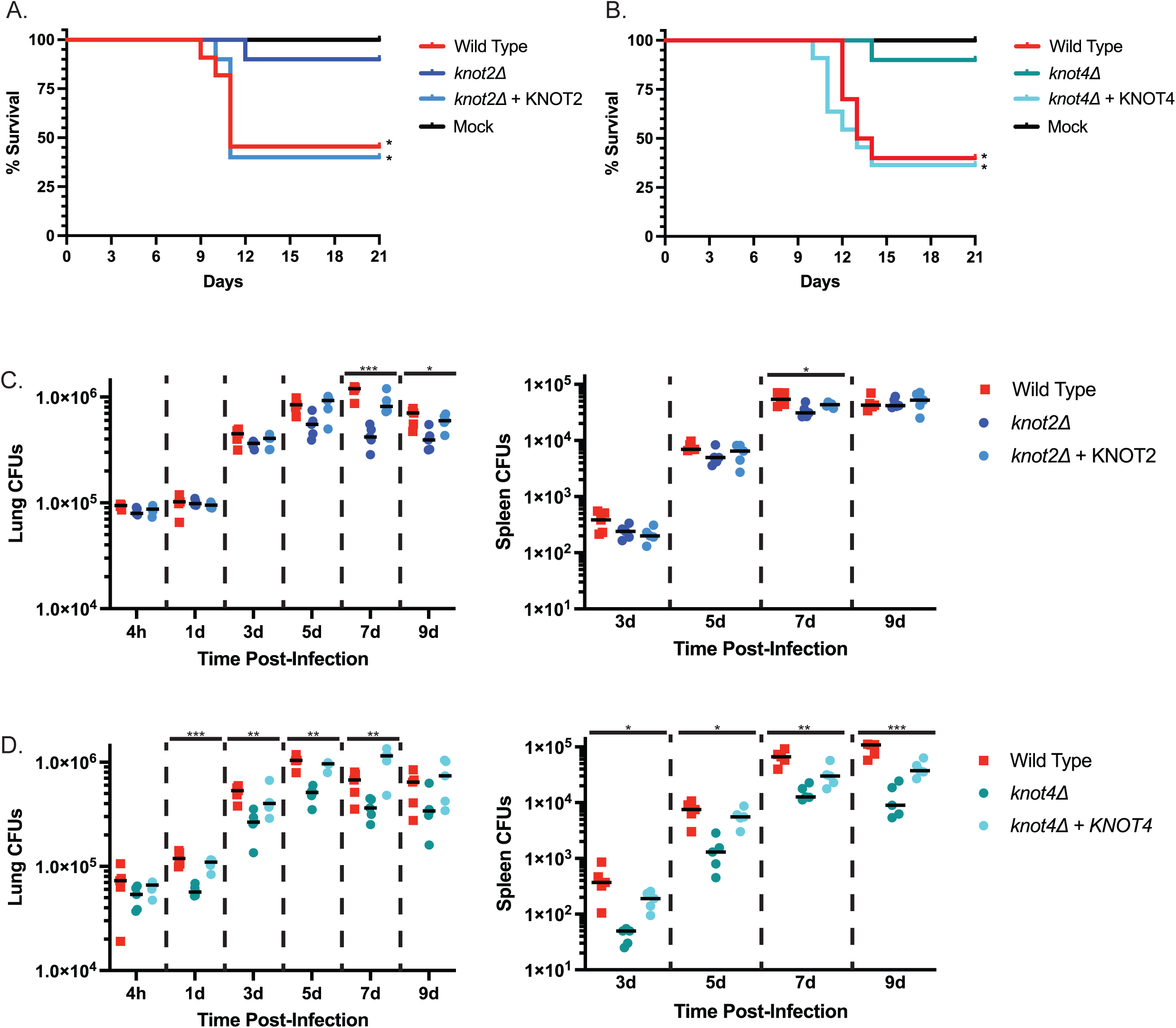
Knot2 and Knot4 are required for virulence *in vivo*. C57BL/6J mice were intranasally infected with 1 x 10^6^ *Histoplasma* yeast (A, B) or 3 x 10^5^ yeast (C, D). (A) Kaplan-Meier survival curve of control vs. *knot2Δ* infection. Mice were mock-infected (n = 3, black), or infected with wild-type (n = 11, red), complement (n = 10, light blue), or *knot2Δ* mutant (n = 10, dark blue) *Histoplasma*. (B) Kaplan-Meier survival curve of control vs. *knot4Δ* infection. Mice were mock-infected (n = 3, black), or infected with wild-type (n = 10, red), complement (n = 11, turquoise), or *knot4Δ* mutant (n = 10, teal) *Histoplasma*. For A and B, asterisks denote p-value < 0.05 according to log rank test. (C) Fungal burden in mouse lungs and spleens for control or the *knot2Δ* mutant (n = 5 per genotype per time point). (D) Fungal burden in mouse lungs and spleens for control or the *knot4Δ* mutant (n = 5 per genotype per time point). For C and D, asterisks denote p-value < 0.05 according to one-way ANOVA test.

Next, we examined whether the *in vivo* virulence defects of the *knot2Δ* and *knot4Δ* mutant strains were due to a reduced ability to colonize the mouse and/or disseminate during infection. C57BL/6J mice were infected intranasally with a sublethal dose of *Histoplasma* (3 x 10^5^ yeast). Mice infected with the *knot2Δ* mutant showed a modest but significant decrease in fungal burden in both the lungs and spleens at seven days post infection (dpi) (Fig. 5C). The *knot4Δ* mutant yeast showed a more significant, persistent decrease in fungal burden in lungs beginning as early as 1 day-post infection (dpi). Fungal burden was also significantly decreased in the spleen starting at 3 dpi, suggesting either a defect in dissemination to the spleen or growth within the spleen (Fig. 5D). Nonetheless, both the *knot2Δ* and *knot4Δ* mutants achieved significant *in vivo* growth. The ability of mice to survive infection with these mutants despite relatively high fungal burden suggests that *KNOT2* and *KNOT4* might influence other processes such as host inflammation.

## Discussion

In this work we apply known characteristic of effectors from plant fungal pathogens to identify putative effectors in the human fungal pathogen *Histoplasma*. Remarkably, we discovered that *Histoplasma* species have undergone a striking expansion of small cysteine knot proteins named knottins. Despite the presence of ∼25 knottin genes per genome, we found that individual knottins are not redundant for pathogenesis—each of the four that we queried is required for virulence in this fungal pathogen of humans.

Many previously identified effectors in plant fungal pathogens tend to be small proteins that are secreted extracellularly and promote host damage and/or cell death by varied mechanisms^11, 46^. Additionally, many have enhanced cysteine content (≥ 4 cysteines), enabling the formation of disulfide bonds that provide stability and rigidity to the protein^47^. Lastly, many of the identified effectors are highly expressed in the infectious form of the fungus^11, 46^, concordant with a role in pathogenesis. We applied these criteria to existing *Histoplasma* datasets, including genomic, transcriptomic and chromatin immunoprecipitation data, successfully identifying 15 putative effectors including three previously identified putative knottins (Fig. 1, S1). As a proof of concept, this set included three proteins (Cbp1, Sod3, and Yps3) that have all been shown to play a role in virulence. Cbp1 is absolutely required for lysis of macrophages and has been shown to induce the integrated stress response (ISR)^34, 36, 37^ to facilitate host-cell death. *Histoplasma* mutants lacking Cbp1 completely fail to kill the host cell but replicate to high levels intracellularly^26^. Intriguingly, Cbp1 has been shown to access the host cytosol through an unknown mechanism^41^. Sod3 is a superoxide dismutase that is required during macrophage infection to protect the yeast against reactive oxygen species (ROS) and is required for *in vivo* virulence^27^. Yps3 is a secreted yeast phase specific protein that localizes to the *Histoplasma* cell wall^28^. Recently, Azimova *et al.* showed that Yps3 binds to Cbp1 and is required for optimal macrophage lysis^41^. Two additional proteins in our set, yeast phase specific protein 21 (YPS21)^57^ and culture filtrate protein 4 (CFP4)^58–60^, have been shown to be preferentially expressed in *Histoplasma* yeast, suggesting a role in pathogenesis. Seven other hits are *Histoplasma* proteins of unknown function with no predicted domains and are fascinating candidates for future analysis. Overall, this computational approach was a robust discovery tool for candidate virulence factors.

Our identification of three knottins in this set of putative effectors is consistent with our original transcriptome annotation identifying a handful of *Histoplasma* knottins that contain a conserved 6-cysteine spacing pattern akin to some insect toxins^55, 85^. Here we wrote a novel algorithm to detect knottins across the *Histoplasma* genome based on this spacing pattern (Fig. 2A, 2B) and the known knottin intron-exon structure^67^. Surprisingly, we identified 25 putative knottins, 13 more than previously reported (Fig. S3). To note, one of these new knottins was in our list of 15 putative effectors (Fig. 1C, S1). Our algorithm provides a robust and sensitive analysis allowing the detection of knottin motifs in previously unannotated regions of the G217B genome^69, 70^ (Fig. S2). Furthermore, genomic locations of the 25 knottins and the identified putative effectors from Figure 1 showed no clustering or preference for any one region of the genome (Fig. S5), giving no insight to the mechanism of knottin expansion in *Histoplasma*. Further bioinformatic analysis identified 22 to 27 knottins across different *Histoplasma* strains (G186AR, H88, H143, WU24 and Tmu), demonstrating that knottin expansion is consistent in these sequenced *Histoplasma* isolates.

Intriguingly, we found few knottins across the fungal kingdom, indicating that the knottin expansion is unique to *Histoplasma* and closely related fungi. On average, most fungal genomes contained fewer than 5 knottins (Fig. 2C). Using published AlphaFold2 structure predictions for 48 proteomes from bacteria to humans, we observed *Histoplasma* has the largest number of predicted knottins among the sampled fungi. The human genome contained 7 predicted knottins of varied function, including knottins involved in appetite regulation and melanin production^86^ (Fig. 2D). Intriguingly, we observed no knottin proteins in prokaryotes, suggesting that a eukaryotic-specific feature like the ER is needed to facilitate knottin folding. Additionally, our analysis indicates that AlphaFold2 is not biased towards false positive knottin fold predictions and is, therefore, a useful tool for discovering new knottins. Likewise, the AlphaFold2 results corroborate that the expansion originally detected by the KNOTTIN_FINDER algorithm is unique and specific to *Histoplasma* genomes. Since *Histoplasma* is an intracellular pathogen, it is compelling to speculate that it requires highly stable knottin proteins to execute virulence-relevant functions in the host cytosol during macrophage infection.

*Histoplasma* knottins display remarkable versatility in their predicted structure. We employed the use of protein structure prediction tools like AlphaFold2 and Modeller to model all 25 putative knottin proteins, along with APBS to predict electrostatic surface potentials. To our surprise, no two knottins were predicted to look exactly alike (Fig. S7). Knottin clades were determined by sequence similarity, and we occasionally observed some pockets of similarity in predicted knottin structure within a clade. For example, clade 11 contains a small stretch of amino acids upstream of the first cysteine of the knottin motif that contains similarity to the RxLR motif that is required for translocation of plant fungal effectors into the host cell^87^ (Fig. S6). It remains to be seen if this motif has biological function in *Histoplasma* knottins. Interestingly, overall similarity between knottin predicted structures was low, which might imply distinct biological functions for individual knottins, or might, as in the case of chemokine-binding tick knottins, reflect the diverse set of binding surfaces necessary to counteract a corresponding defensive diversity in unknown host targets^88^. Future analysis of knottin function will shed light on the need for such a large plethora of knottins in *Histoplasma* biology and reveal whether knottins modulate conserved processes in the host. Additionally, it will be fascinating to understand if *C. fulvum AVR9*, the founding member of the knottin Fungi1 family, and particular *Histoplasma* knottins such as its homolog *KNOT4*, target similar processes in plant and animal cells.

To validate our computational approach and explore the role of knottins in *Histoplasma* pathogenesis directly, we selected *KNOT1-4* for further characterization. *KNOT1*, *KNOT2*, and *KNOT3* were interesting to us since they were identified in our initial analysis of putative effectors (Fig. 1, S1) and *KNOT4*, as mentioned above, is the homolog of *C. fulvum AVR9*. Furthermore, *KNOT1-4* are from distinct gene clades, providing a broad initial interrogation of the role of key knottins in pathogenesis (Fig. S6, S7). Of note, *KNOT1* and *KNOT4* are each unique within the G217B genome. In contrast, *KNOT2* has one paralog and *KNOT3* has six (Fig. S9), which could suggest functional redundancy between some of the knottins. Despite the plethora of *KNOT3* paralogs, we observed that *Histoplasma* yeast lacking *KNOT3* had a modest growth defect *in vitro* (Fig. S11) that was exacerbated in macrophages (Fig. 3), suggesting that Knot3 may play a general role in *Histoplasma* growth even outside of the host cell. For example, it could be that Knot3 plays a role in nutrient acquisition that becomes even more salient within the macrophage phagosome, where iron, zinc, nucleic acids and vitamins are limiting^89^. *KNOT1*, *KNOT3,* and *KNOT4* were each required for macrophage lysis and intracellular growth whereas *KNOT2* was dispensable for intracellular growth but required for macrophage lysis (Fig. 3). Intriguingly, host-cell death was significantly reduced in these mutants but not abrogated, suggesting either that other knottins or other effectors contribute to cell death.

We observed that Knot1-4 proteins were present in the macrophage cytosol during infection (Fig. 4B), much like the virulence factor Cbp1^41^. These data suggest that these effectors have targets within the host cell. The mechanism of how *Histoplasma* effectors gain access to the macrophage cytosol is unknown. Interestingly, Cbp1 is required to trigger the host ISR, but individual knottins are not, despite their partial macrophage lysis defect (Fig. 4C). These data suggest that host pathways other than the ISR must be involved in knottin-mediated cell lysis, or that we cannot detect small changes in ISR activation that could be relevant to individual knottin mutants. Future studies will examine the consequences of deleting more than one knottin gene on ISR activation and macrophage lysis.

The large number of knottins proteins could not compensate for the lack of either *KNOT2* or *KNOT4* in mice; these knottins were required for wild-type virulence of *Histoplasma* (Fig. 5A, B). Importantly, mice infected with these knottin mutants experienced weight loss (Fig. S12), indicating that the mice manifested symptoms of illness but most were able to recover. *KNOT2* was largely dispensable for fungal burden in mice, except for a modest decrease in both the lungs and spleens at seven days post infection (Fig. 5C). These data are concordant with the role of Knot2 in macrophage infections, since we observed that Knot2 is dispensable for intracellular growth but required for macrophage lysis. In the case of *KNOT4*, the mutant displayed reproducibly lower fungal burden in mice at all time points examined in both lung and spleen, suggesting a modest defect in fungal dissemination and/or replication (Fig. 5D). These data contrast with the phenotype of the *knot4Δ* mutant in macrophage infection, where there seems to be a more significant effect on intracellular fungal burden. Overall, although there is a statistically significant decrease in fungal burden during infection with each mutant when compared to wild-type, the mutants are still present at relatively high levels and fail to be cleared by the host. The fact that Knot2 and Knot4 are largely dispensable for fungal burden in the mouse but are required for *Histoplasma* to kill the host efficiently suggests that they may affect processes independent of or in addition to fungal replication. Ultimately, a full understanding of knottin biology in *Histoplasma* pathogenesis will include generating an atlas of host responses to a variety of knottin mutants, including their impact on cytokine production and host inflammation. What is clear is that this family of intriguing effector proteins plays key roles in *Histoplasma* pathogenesis.

## Acknowledgements

This research was supported by NIH 1R01AI172258-01A1 (to AS) and NIH R37AI066224 (to AS) for funding. AS is a Chan Zuckerberg Biohub – San Francisco Investigator. We acknowledge the UCSF CAT for sequencing resources. The funders had no role in study design, data collection and interpretation, or the decision to submit the work for publication.

## Materials and Methods

### KNOTTIN_FINDER

Knottins in fungal genomes were predicted by running KNOTTIN_FINDER (Supp code). Specifically, KNOTTIN_FINDER scans all six reading frames to find open reading frames (i.e. subsequences not containing stop codons) of < 750 nucleotides with at least 6 cysteines and matching the spacing pattern: CX3-4CX5CX3-6CX2CX6-12C. ORFs matching this pattern were removed if they overlapped annotated genes with CDS ≥ 750 nucleotides to select for small proteins.

### Phylogenetic tree analysis

The KNOTTIN_FINDER algorithm was run against 56 select fungal genomes (Table 1). Putative knottins were aligned to the Fungi1 HMM (hidden Markov model) from Gilmore *et al.* with hmmalign from HMMer 3.3.2^90^ and a gene tree was generated from this alignment using FASTTREE 2.1.11^91^. A corresponding species tree was generated by aligning the monophyletic Gcd10p family to the Pfam Gcd10p HMM with hmmalign and generating a phylogeny with FASTTREE. Clades were manually annotated in the knottin tree and genes per clade (horizontal) were plotted against the Gcd10p-based species tree (vertical) as a heatmap. Final array tree branch lengths are not to scale.

### Protein alignments

Individual probcons (Probabilistic Consistency-based Multiple Alignment of Amino Acid Sequences)^92^ alignments of knottins clades were run to identify paralogs within the different *Histoplasma* species (G217B, G186AR, WU24, H88, H143 and Tmu). Clustal omega^93^ alignments of the C-terminal region of knottins were run against 25 putative *Histoplasma* knottins. Alignments were manually organized numerically by clade. Amino acids were colored based on physiochemical properties.

### KNOTTIN_TOPOLOGY_SCAN

Tarballs of AlphaFold2 predictions for 48 proteomes^77^ (Table 2) were downloaded from https://alphafold.ebi.ac.uk/ from 8/21/2023 - 8/24/2023. KNOTTIN_TOPOLOGY_SCAN was used to detect the knottin fold in each structure as follows (Data S2, Fig. S8). Cysteines within 3 Angstroms of each other were considered disulfide bonded. As shown in Figure S8, for each set of six cysteines with the C1-C4, C2-C5, C3-C6 disulfide bonding pattern, we defined the embedded ring as the covalent bonds from the alpha carbon of C1, through the backbone to C2 (yellow), through the C2-C5 disulfide to C5 (green), through the backbone to C4 (orange), and through the C4-C1 disulfide back to the alpha carbon of C1 (red). Embedded rings longer than 26 residues were discarded, based on the largest embedded ring observed in KNOTTINDB. The ring surface (grey) was defined as the set of triangles formed by connecting each edge of the ring with the centroid of the ring. The locking element was defined as the covalent bonds from the alpha carbon of C2 through the backbone to C3 (cyan), through the C3-C6 disulfide to C6 (blue), through the backbone to C5 (magenta). A set of six cysteines was considered to have the knottin fold if the locking element made an odd number of crossings through the ring surface.

### Alphafold predictions

Mature sequences (includes knottin motif and three amino acids preceding the first cysteine) for the 25 predicted knottins from KNOTTIN_FINDER for *Histoplasma* G217B were used for predictions. Predicted structures for these sequences were generated by running AlphaFold v2.3.0^94^ on the UCSF Wynton HPC cluster.

### MODELLER predictions

The mature protein sequences for the 25 *Histoplasma* G217B knottins detected by KNOTTIN_FINDER were aligned with PROBCONS 1.12^92^. The resulting alignment was converted to PIR format and linked to the AlphaFold2 predicted structures for the 17 sequences where the top ranked prediction had a knottin fold. Homology models for each of the 25 mature sequences were then generated by MODELLER 9.17^78^ in automodel mode using all 17 structures as templates.

### Electrostatic potential surfaces

Electrostatic potentials for the MODELLER predicted structures were calculated using PDB2PQR 3.5.2 and APBS 3.4.1^79^ and potentials at the molecular surface were rendered using PYMOL 2.5.0 with post-processing in IMAGEMAGICK 6.9.11. Consistent orientation of knottins for rendering was achieved by rigid body alignment of the cysteine C alpha atoms to a reference orientation using the PYMOL fit command.

### Generation of Histoplasma strains

*Histoplasma* strain G217B *ura5Δ* (WU15) was a kind gift from the lab of William Goldman (University of North Carolina, Chapel Hill). For all studies in this paper, “wild-type” refers to G217B *ura5Δ* transformed with a *URA5*-containing episomal plasmid. The knottin deletion mutants were all generated from the G217B *ura5Δ* parental strain transformed with an episomal plasmid which contains a bi-directional H2AB promoter driving Cas9 and two single guide RNA cassettes targeting sequences at the start and end of G217B knottin genes (*KNOT1-4*). Single colony isolates were screened by PCR to identify edited genomic sites. Single colonies were repeatedly isolated and screened until the wild-type copy of the gene of interest was not detectable by PCR. Plasmid containing the Cas9 was lost from the mutants by growing the *Histoplasma* yeast in HMM (*Histoplasma* macrophage media) containing exogenous uracil and then colonies were screened for plasmid loss. The purified mutant *Histoplasma* yeast (*ura5Δ knot1Δ*, *ura5Δ knot2Δ, ura5Δ knot3Δ, or ura5Δ knot4Δ)* were transformed with either the *URA5-*containing episomal plasmid or a URA5-containing complementation vector. Complementation constructs contained native promoters (including both upstream and downstream UTRs) and any binding sites for the Ryp transcription factors. All tagged knottin strains used in this study including Knot1-3XFLAG, Knot2-3XFLAG, Knot3-3XFLAG and Knot4-3XFLAG were C-terminally 3X-FLAG tagged under the control of their native promoters and constructs were introduced into *Histoplasma* G217B *ura5Δ* strain.

### Concentrating Histoplasma culture supernatants

*Histoplasma* yeast were grown in HMM at 37°C, 5% CO_2_ in an orbital shaker until stationary phase. 10 mL of culture were spun down at 2500 rpm for 5 minutes. Supernatants were passed through a 0.22 μm filter to sterilize. Sterile supernatants were concentrated using MilliporeSigma^TM^ Amicon^TM^ centrifugal filter units with a 3 kDa molecular weight cut off (MWCO) to preserve small proteins. Culture supernatants were concentrated ∼100-fold, snap frozen, and stored at −80°C until ready to use.

### BMDM differentiation and culture conditions

Bone marrow derived macrophages (BMDMs) were isolated from the tibias and femurs of 6-8 week-old C57BL/6J (Jackson Laboratories stock, No. 000664) mice. Mice were euthanized via CO_2_ narcosis and cervical dislocation as approved under UCSF Institutional Animal Care and Use Committee protocol. Cells were differentiated in BMM (bone marrow macrophage media) containing Dulbecco’s Modified Eagle Medium (DMEM High Glucose), 20% Fetal Bovine Serum, 10% v/v CMG supernatant (the source of CSF-1), 2 mM glutamine, 110 μg/mL sodium pyruvate, 110 μg/mL penicillin and streptomycin. Undifferentiated monocytes were plated in BMM for 7 days at 37°C and 5% CO_2_. Adherent cells were collected and frozen in 40% FBS and 10% DMSO and stored in liquid nitrogen for future use.

### Macrophage infections

Macrophage infections with G217B *Histoplasma* strains were performed as described previously^26, 30, 37, 41^. Briefly, the day before infection, macrophages were seeded in tissue culture-treated dishes in triplicate. On the day of infection, yeast cells from logarithmic-phase cultures (OD_600_ = 4-7) were collected, resuspended in D-PBS (PBS Ca^2+^, Mg^2+^ free), sonicated for 3 seconds on setting 2 using a Fisher Scientific Sonic Dismembrator Model 100, and counted using a hemacytometer. After a 2-hour phagocytosis period, the macrophages were washed with D-PBS to remove any extracellular yeast and fresh BMM media was added. For infections lasting longer than 2 days, fresh BMM media was added to the cells. As a control for ISR induction, 2.5 μg/mL final concentration of Tunicamycin (Santa Cruz Biotechnology) was added to cells.

### Lactate dehydrogenase release assay

To quantify macrophage lysis, BMDMs were seeded (7.5 x 10^4^ cells per well of a 48-well plate) and infected in triplicate as described above. At the indicated time points, the amount of LDH in the supernatant was measured as described previously^95, 96^. BMDM lysis was calculated as the percentage of total LDH from supernatant of wells with uninfected macrophages lysed in 1% Triton X-100 at the time of infection. Due to continued replication of BMDMs over the course of the experiment, the total LDH at later time points can be greater than the total LDH from the initial time point, resulting in an apparent lysis that is greater than 100%.

### Intracellular replication

BMDMs were seeded and infected in triplicate as described above. At the indicated timepoints, culture supernatants were removed and ddH_2_O was added to cells. Cells were incubated at 37°C for 15 min and then mechanically lysed by vigorously pipetting. The lysate was collected, sonicated to disperse any yeast clumps, and yeast cells were counted on a hemocytometer. Appropriate dilutions were performed, and *Histoplasma* was plated on HMM agarose plates. Plates were incubated at 37°C with 5% CO_2_ for 12-14 days before CFUs were enumerated. To prevent any extracellular replication from confounding the results, intracellular replication was not monitored after the onset of macrophage lysis.

### Fractionation of Histoplasma-infected macrophage lysates

BMDMs (2 x 10^7^ cells) were plated on 15 cm TC-treated plates and infected with different *Histoplasma* strains at MOI of 5 yeast to 1 macrophage. At 24 hours post infection, cells were gently collected by scraping without washing. Cells were spun down at 2500 rpm for 5 min to pellet at 4°C. The cell pellet was resuspended in 500 μL of D-PBS (Ca^2+^, Mg^2+^ free) and transferred to a 1.5 mL eppendorf tube and spun at 1000 rpm for 2 min to wash cells. 300 μL of homogenization buffer (20 mM HEPES pH 7.4, 150 mM KCl, 2 mM EDTA, cOmplete Mini Protease Inhibitor Cocktail tablet-Roche) was used to resuspend cell pellets. The cell lysate was passed through a 27-gauge needle to ensure only the plasma membrane was ruptured, leaving internal membranous organelles intact. The lysate was spun at 3000 rpm for 5 minutes to pellet *Histoplasma* cells and macrophage nuclei. The remaining lysate was spun at 45,000 rpm for 2 hours in a TL-100 tabletop ultracentrifuge. The resulting supernatants representing the cytosolic fraction were transferred into 1.5 mL low-binding tubes. The pellet (membrane fraction) was resuspended in homogenization buffer with 1% Triton X-100. All samples were flash frozen in liquid nitrogen and stored at −80°C.

### DNA isolation and PCR

DNA was extracted from two-day late log *Histoplasma* cultures using the Qiagen Gentra Puregene Yeast/Bact. Kit according to manufacturer protocol. Takara PrimeSTAR GXL DNA polymerase was used to amplify genomic regions per manufacturer protocol. Primers used are listed in Table 3. PCR products were run on a 1% agarose gel and imaged on a Bio-Rad Gel Doc^TM^ EZ Imager.

**Table 3:**
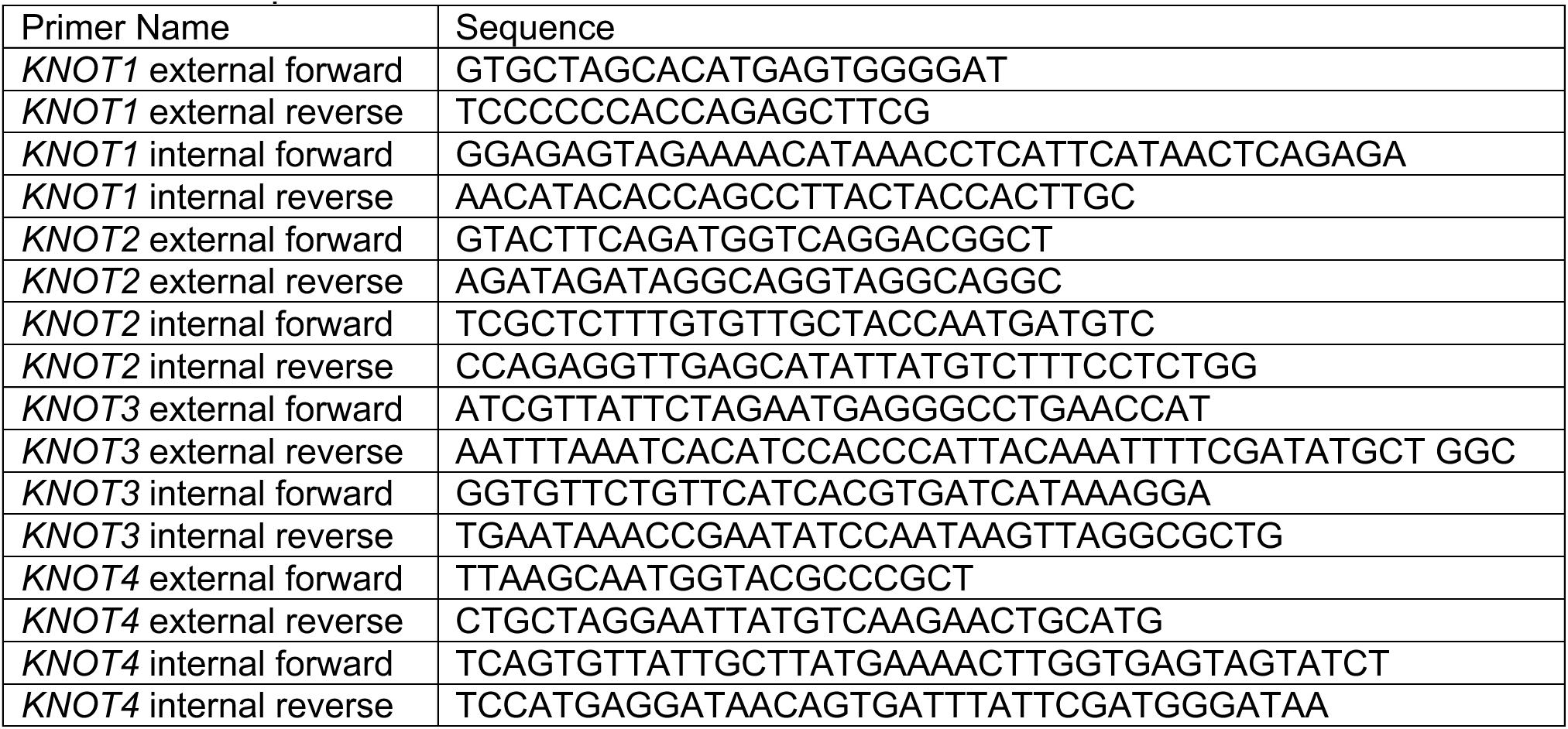
List of primers used to screen *knot1-4Δ* CRISPR deletion mutants.

### Whole genome sequencing

250 ng of genomic DNA (purified as described above) was used to generate libraries using Nextera DNA prep library kit as per manufacturer protocol. Individual libraries were uniquely barcoded using Illumina’s Nextera DNA Indexes (i5 and i7). The final libraries were sequenced on Illumina NovaSeq600 sequencer.

### RNA isolation and protein extraction

10 mL of two-day late log *Histoplasma* yeast cultures were pelleted and resuspended in 1 mL of QIAzol (Qiagen). Samples were bead-beaten in a Mini-BeadBeater-96 (BioSpec 1001) with zirconia beads for 2 minutes to lyse cells. Chloroform (200 μL) was added and vigorously vortexed for 15 seconds. Lysates were centrifuged at 12,000xg for 20 minutes at 4°C. For RNA isolation from cultured cells, infected macrophages were lysed in 1 mL total of QIAzol (Qiagen) Total RNA was isolated from the aqueous phase using RNA columns (Epoch Life Science) and then subjected to on-column DNase digestion (Invitrogen PureLink). Total RNA was eluted using nuclease free water. For protein extraction, pure ethanol was added to organic phase and centrifuged at 2000xg for 5 minutes to pellet DNA. Isopropanol was added and samples were centrifuged at 12,000xg to pellet proteins. Protein pellets were washed with 0.3 M guanidine thiocyanate followed by 100% ethanol wash. Protein pellets were resuspended in urea lysis buffer (9 M urea, 25 mM Tris-HCl pH 6.8, 1 mM EDTA pH 8.0, 1% SDS, and 0.7 M βME).

### qRT-PCR

For cDNA synthesis, 2-4 μg total RNA was reverse transcribed using Maxima H Minus Reverse Transcriptase (Fisher Scientific), oligo-dT and pdN9 oligos following manufacturer’s protocol. Quantitative PCR was performed on 1:10 to 1:50 dilutions of cDNA template using FastStart Universal SYBR Green Master Mix (Rox) (Sigma-Aldrich). Reactions were run on an Mx3000P machine (Agilent) and analyzed using MxPro software. Abundances of mouse *TRIB3* were normalized to mouse HPRT levels. *KNOT1*, *KNOT2*, *KNOT3* and *KNOT4* levels were normalized to *Histoplasma GAPDH* levels. Primer sequences are listed in Table 4.

**Table 4:**
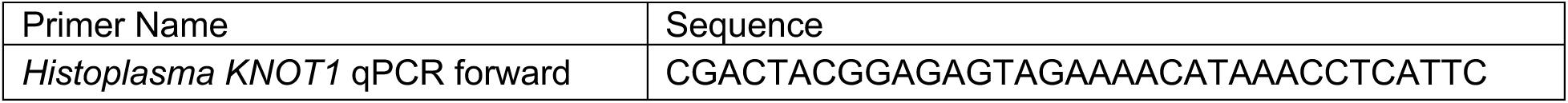
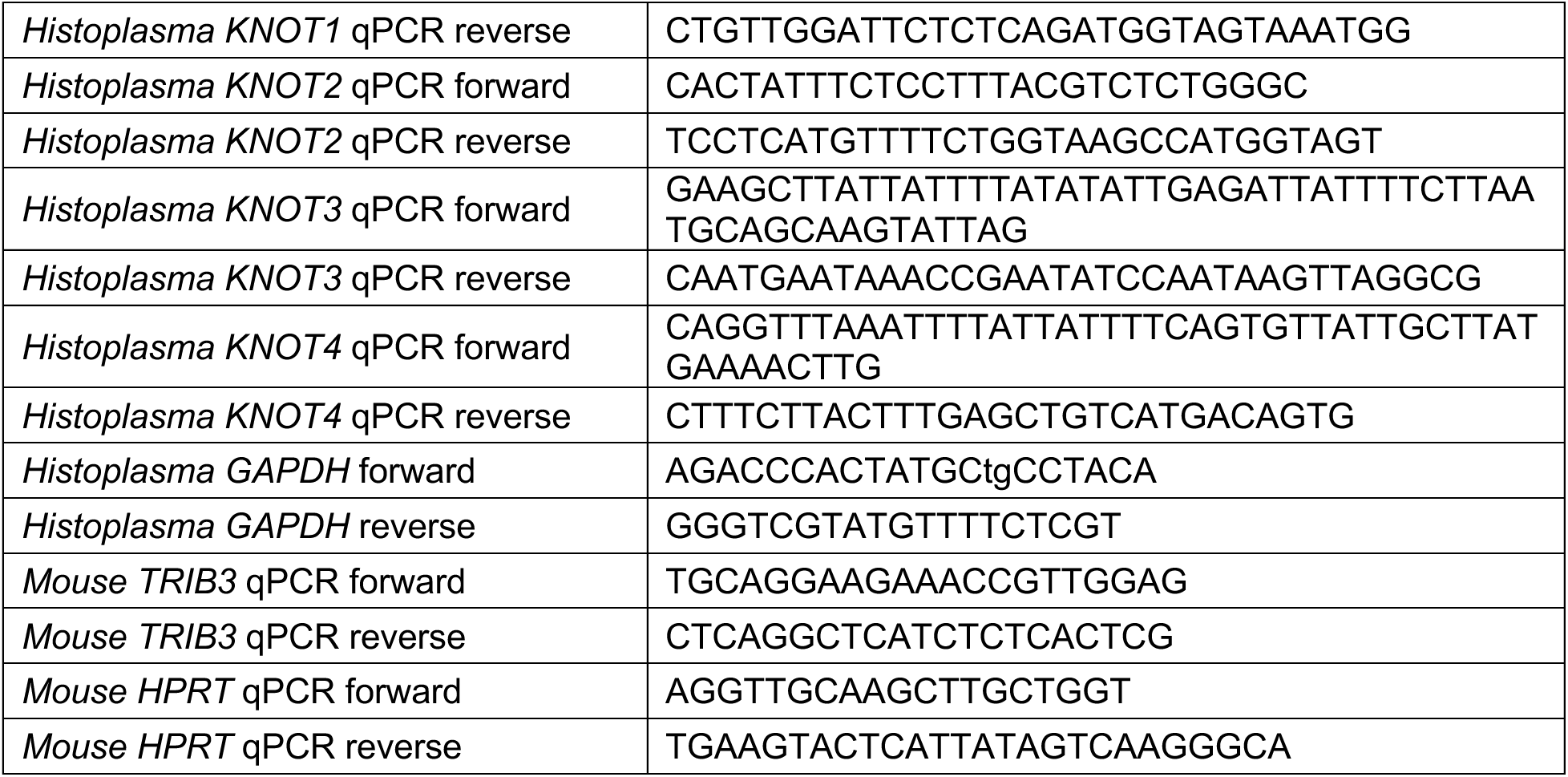
List of primers used for qRT-PCR analysis for expression in *Histoplasma* and *TRIB3* expression in macrophages.

### SDS-PAGE Protein gel and western blot analysis

Protein samples were mixed with protein loading buffer (LI-COR for fractionation experiments or NuPAGE for yeast samples), 50mM DTT, and denatured at 95°C for 5 minutes. Proteins were separated by SDS-PAGE on NOVEX-NuPAGE 4-12% BIS-TRIS gels using 20X NuPAGE MES Running Buffer (Invitrogen). Precision Plus Dual Xtra Protein Standards (Bio-Rad) were used to estimate the molecular weight of proteins. For Western blots, the SDS-PAGE separated proteins were transferred to nitrocellulose membranes. Non-specific binding to the membrane was blocked with LI-COR Intercept (PBS) blocking buffer and probed with the antibodies listed below. Blots were imaged on an Odyssey CLx and analyzed using ImageStudio2.1 (LI-COR). The following primary antibodies were used: monoclonal mouse anti-FLAG M2 (Millipore Sigma F3165), custom generated rabbit anti-Cbp1^41^, custom generated rabbit anti-Ryp1^56^, mouse anti-ɑ-tubulin (DM1A) (Novus biological NB100-690), and rabbit anti-calnexin (abcam 22595).

### In vitro growth

Two-day, late-log *Histoplasma* cultures were used to inoculate 100 mL of HMM medium to a starting OD_600_ = 0.05 in triplicate. Cultures were incubated in an orbital shaker at 37°C with 5% CO_2_ for 144 hours. At each timepoint, 1 mL of each culture was removed using a serological pipette, vigorously vortexed for 30 seconds, diluted to be within the linear range, and analyzed to determine OD_600_ on spectrophotometer (Eppendorf BioPhotometer).

### Mouse infections

Eight- to twelve-week-old female C57BL/6J mice (Jackson Laboratory stock, No. 000664) were anesthetized with isoflurane and infected intranasally with wild-type *Histoplasma* (G217B *ura5Δ* + URA5), the *knot2Δ* mutant (*knot2Δ* + URA5), the complemented stain (*knot2Δ* + KNOT2), the *knot4Δ* mutant (*knot4Δ* + URA5) or the complemented stain (*knot4Δ* + KNOT4). The inoculum was prepared by collecting mid-logarithmic phase (OD_600_ = 4-7) yeast cultures, washing once with D-PBS (Ca^2+^, Mg^2+^ free), sonicating for 3 seconds on setting 2 using a Fisher Scientific Sonic Dismembrator Model 100, counting with a hemacytometer, and diluting with D-PBS so the final inoculum was approximately 30 μL total. To monitor survival, animals were infected intranasally with a lethal dose of 1 x 10^6^ yeast per mouse. For survival curve analysis, mice were euthanized after they exhibited three days of sustained weight loss greater than 25% of their initial weight in addition to one other disease symptom (hunching, panting, ears tucked back, and/or lack of grooming). To monitor fungal burden, animals were infected intranasally with a sublethal dose of 3 x 10^5^ yeast per mouse. At every time point, 5 mice per genotype were euthanized and lungs and spleens were harvest. Organs were homogenized in D-PBS, serially diluted and plated on brain heart infusion (BHI) agar plates supplemented with 10% sheep’s blood. CFU’s were enumerated after 10–12 days of growth at 30°C.

## Data availability statement

The raw sequencing data are available at the NCBI Sequence Read Archive (SRA) under SRA accession PRJNA1199544.

