## Supplemental figures for "A remarkable expansion of secreted cysteine knot proteins reveals novel virulence factors in the human fungal pathogen *Histoplasma*"

**Supplemental Figure 1. Expression pattern of the 15 putative *Histoplasma* virulence factors.**

In (1), ribosome occupancy on transcripts preferentially expressed in the yeast form (red) from published ribosome profiling data[55] is shown. Y-axis is scaled to max signal for gene of interest. In (2), expression of yeast transcripts (red) from published transcriptomics data of steady-state yeast[55] is shown. Y-axis is scaled to max signal for gene of interest. In (3), paired-end (PE) transcript model of genes in *Histoplasma* (CDS light gray, UTR dark gray) is shown. Vertical lines on *KNOTTIN_006* (*KNOT2*) (blue), *KNOTTIN_009* (*KNOT1)* (orange), and *KNOTTIN_023* (*KNOT3)* (purple) denote Cas9 guide RNA locations used for generating deletion strains used in this study. In (4), smoothed ChIP-chip signal[56] (red = Ryp1, green = Ryp2, blue = Ryp3, purple = Ryp4) is shown. Y-axis is scaled to chip enrichment at max 10 for all genes.

**Supplemental Figure 2. KNOTTIN_FINDER algorithm identifies putative knottin genes in unannotated genomic regions.**

Genome browsers of two representative knottins showing assembled transcripts based on PE transcriptomic data, ribosome occupancy from steady state yeast cultures, and predicted transcripts. (A) Putative knottin denoted by a blue line identified in a previously annotated gene. (B) A putative knottin (blue line) in a previously unannotated region. Ribosome occupancy is observed where the knottin motif is denoted, indicating that translation occurs. Triangles (red and blue) denote coding sequence in both the assembled and predicted transcripts.

**Supplemental Figure 3. Ribosome profiling and transcriptomics results for the remaining 21 putative knottins.**

*KNOTTIN_006, 009, 022, 023* are excluded as they are shown in Figure S1. In (1), ribosome occupancy on transcripts preferentially expressed in the yeast form (red) from published ribosome profiling data[55] is shown. Y-axis is scaled to max signal for gene of interest. In (2), expression of yeast transcripts (red) from published transcriptomics data of steady-state yeast[55] is shown. Y-axis is scaled to max signal for gene of interest. In (3), paired-end (PE) transcript model of genes in *Histoplasma* (CDS light gray, UTR dark gray) is shown where available. Horizontal blue lines denote the location of each knottin motif. On *KNOTTIN_025* (*KNOT4*), the homolog of *C. fulvum AVR9*, vertical teal lines denote Cas9 guide RNA locations used for generating the deletion strain used in this study.

**Supplemental Figure 4: Effector criteria applied to the 25 putative knottins.**

Six knottins (*KNOTTIN_002, 003, 007, 008, 011,* and *018*) are not part of this analysis due to their absence in the PE genome assembly. Four knottins (*KNOTTIN_004, 019, 020*, and *021*) were identified to be small, cysteine-rich, and yeast-enriched. Eleven knottins (*KNOTTIN_000, 001, 005, 010, 012, 013, 014, 015, 016, 017,* and *025*) were identified as being small, secreted, cysteine-rich, and yeast-expressed. Finally, four knottins (*KNOTTIN_006, 009, 022,* and *023*) are found to be small, secreted, cysteine-rich, Ryp targets and yeast-enriched. These four were identified in Figure 1 and plots for these knottins are shown in Figure S1.

**Supplemental Figure 5. Nanopore sequencing reveals random distribution of knottins and putative virulence factors throughout the G217B *Histoplasma* genome.**

Chromosome map reveals knottins are randomly distributed across the seven chromosomes of the *Histoplasma* G217B genome. Knottin locations are marked by red vertical lines below the chromosomes (black horizonal lines). Repeat regions such as transposons are denoted by purple blocks across the chromosomes. The 15 putative effectors identified in Figure 1 are marked by black vertical lines and names above the chromosomes.

**Supplemental Figure 6. Protein alignment of 25 putative knottins.**

Clustal Omega protein alignment of the 25 putative knottins C-terminal regions (containing the knottin motif plus 20 amino acids upstream of the first cysteine in the motif). Cysteines in the knottin motif are denoted in yellow. Consensus of any amino acids at the different positions across this region are denoted above. Areas of high conservation are highlighted by colors depending on the properties of the amino acids. Hydrophobic residues are in pink, hydrophilic residues are highlighted in purple, positive and negative amino acids are highlighted in red and blue respectively. Aromatic amino acids are marked in green and conformationally special amino acids are highlighted in orange. Knottins that are further characterized in this study are highlighted in grey. Knottins are listed by clade in numerical order. Conservation at each amino acid position is denoted by black bars on top.

**Supplemental Figure 7. Predicted protein structures of all putative knottins.**

(A) Predicted protein structures of all putative knottins by MODELLER from AlphaFold2 templates. Colors correspond to disulfides and loops shown in Figure 2A. Pink text refers to structures where AlphaFold2 failed to predict the knottin fold. (B) Electrostatic surface potentials for all 25 putative knottins using APBS are shown. Front and back (180° rotation) renders are shown side by side in A and B. Putative knottins are listed numerically by clade. Knottins further characterized in this study are highlighted by a grey box in A and B.

**Supplemental Figure 8. Visual representation of the KNOTTIN_TOPOLOGY_SCAN algorithm.**

Representative knottin fold is shown with colors that correspond to disulfides and loops shown in Figure 1A. Cysteines are labeled (C1-C6). The embedded ring that is characteristic of a knottin fold is shown in grey. The ring surface was defined as set of triangles as shown.

**Supplemental Figure 9: Knot1-4 protein alignment across different *Histoplasma* species.**

Individual probcons alignments of C-terminal region of Knot1 (Knottin_009), Knot2 (Knottin_006), Knot3 (Knottin_023), and Knot4 (Knottin_025) across different *Histoplasma* species (G186AR, H88, H143, WU24 and Tmu). Knottins are listed numerically by clade. Only one copy of Knot1 and Knot4 are found in any *Histoplasma* species analyzed. Knot2 has one paralog in G217B (Knottin_012) while Knot3 has six paralogs in G217B (Knottin_001, 008, 010, 016, 018, and 021). Paralogs are highlighted in bold text. Knottins characterized further in this study are denoted by a black arrow. Colors correspond to properties of amino acids described in Figure S6.

**Supplemental Figure 10. Deletion and complementation of *KNOT1-4* and validation of strains.**

(A) PCR strategy to identify deletion mutants. Primers were designed to bind outside the predicted cut sites (external primers); bind outside and inside a cut site and amplify across one of the cut sites (external/internal); and amplify inside the genomic region that has been excised (internal primers). Primers used are listed in Table 3. (B) 1% agarose gels stained with ethidium bromide showed PCR results validating deletion mutants from genomic DNA that were generated and used in this study. (C) Genomic DNA from *knot1-4Δ* mutant yeast was extracted and subjected to whole genome sequencing (WGS). Paired-end (PE) transcript model (top) for each knottin is shown (CDS light gray, UTR dark gray). Reads for each deletion mutant strain (middle) and wild-type strain (bottom) were aligned to the G217B reference genome. (D) qRT-PCR further validated the deletion of *KNOT1-4* and confirmed complementation. Expression of genes of interest were all normalized to the housekeeping gene GAPDH. ND = not detected and NS = not significant. Asterisks indicate $p < 0.05$ relative to wild-type according to t-test.

**Supplemental Figure 11. *In vitro* growth curves reveal Knot3 is required for optimal growth.**

Control and *knot1Δ* (A), *knot2Δ* (B), *knot3Δ* (C), or *knot4Δ* (D) strains were grown under standard culture conditions over 144 hours. Asterisks indicate $p < 0.05$ relative to wild-type according to t-test.

**Supplemental Figure 12. Mice infected with *knot2Δ* or *knot4Δ* mutants lose weight but do not succumb to infection.**

Mice were infected with control or mutant strains at a dose of $1 \times 10^6$ yeast. Average weight of mice was normalized to day 0 weight over the course of 21 days. Mice infected with the *knot2Δ* mutant (A) are shown with dark blue squares whereas mice infected with the *knot4Δ* mutant (B) are shown with teal squares. 25% weight loss (euthanasia criterion) is denoted by the black dashed line.

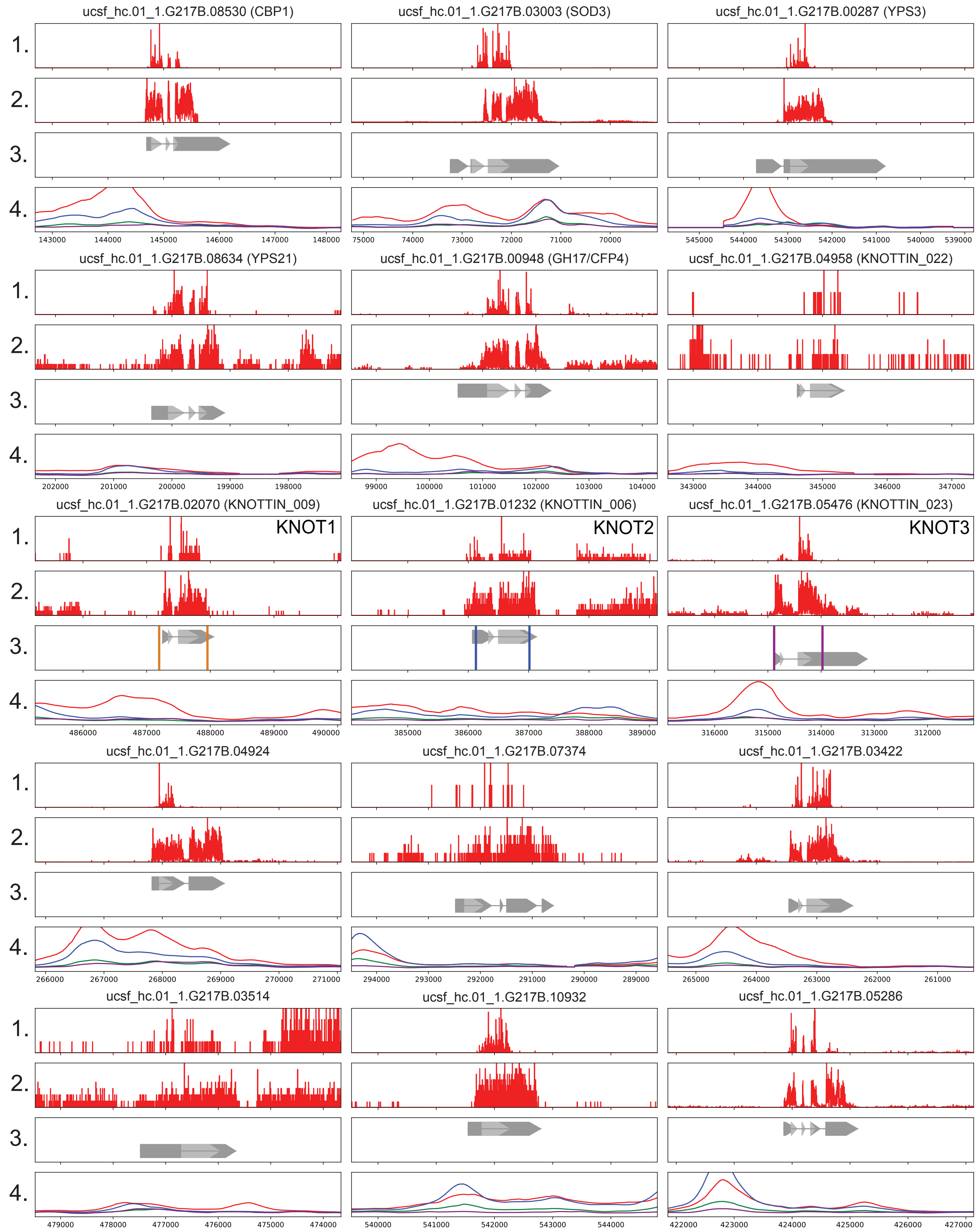


Figure S1. Expression of the 15 putative *Histoplasma* virulence factors.

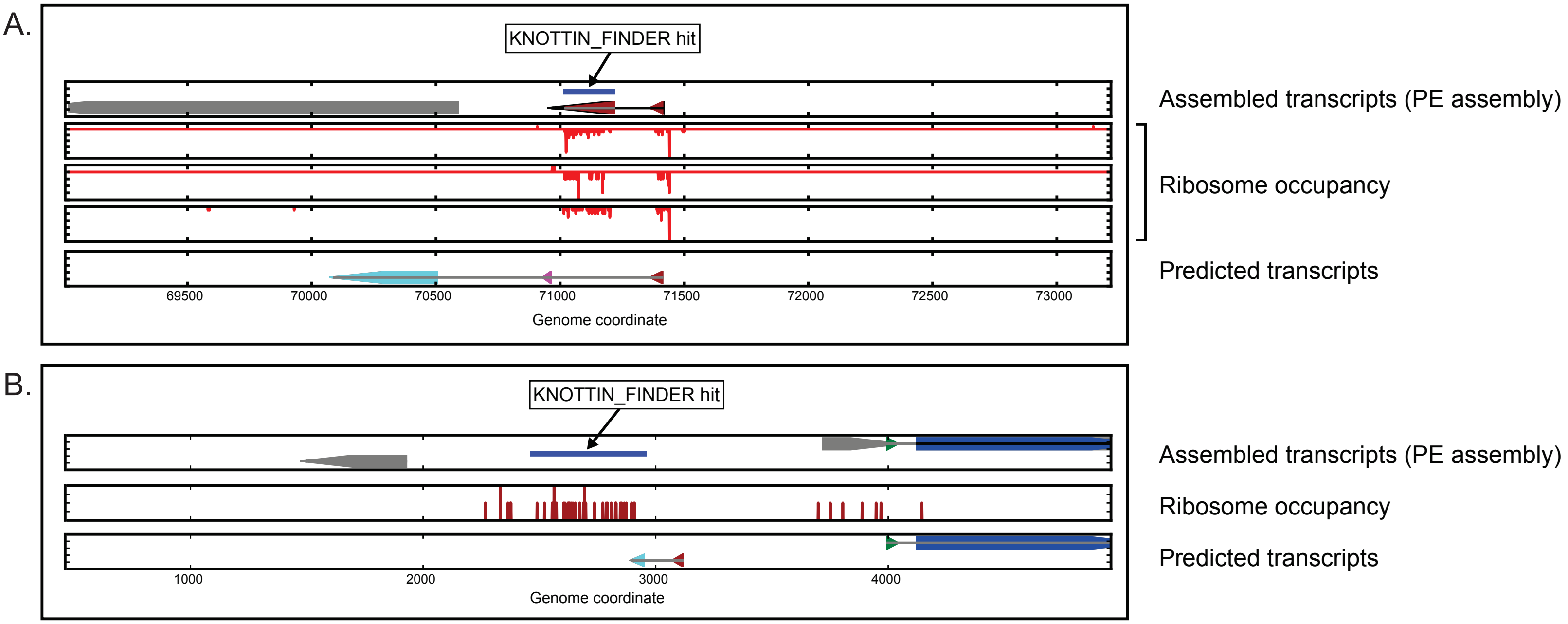


Figure S2. KNOTTIN_FINDER algorithm identifies putative knottin genes in unannotated genomic regions.

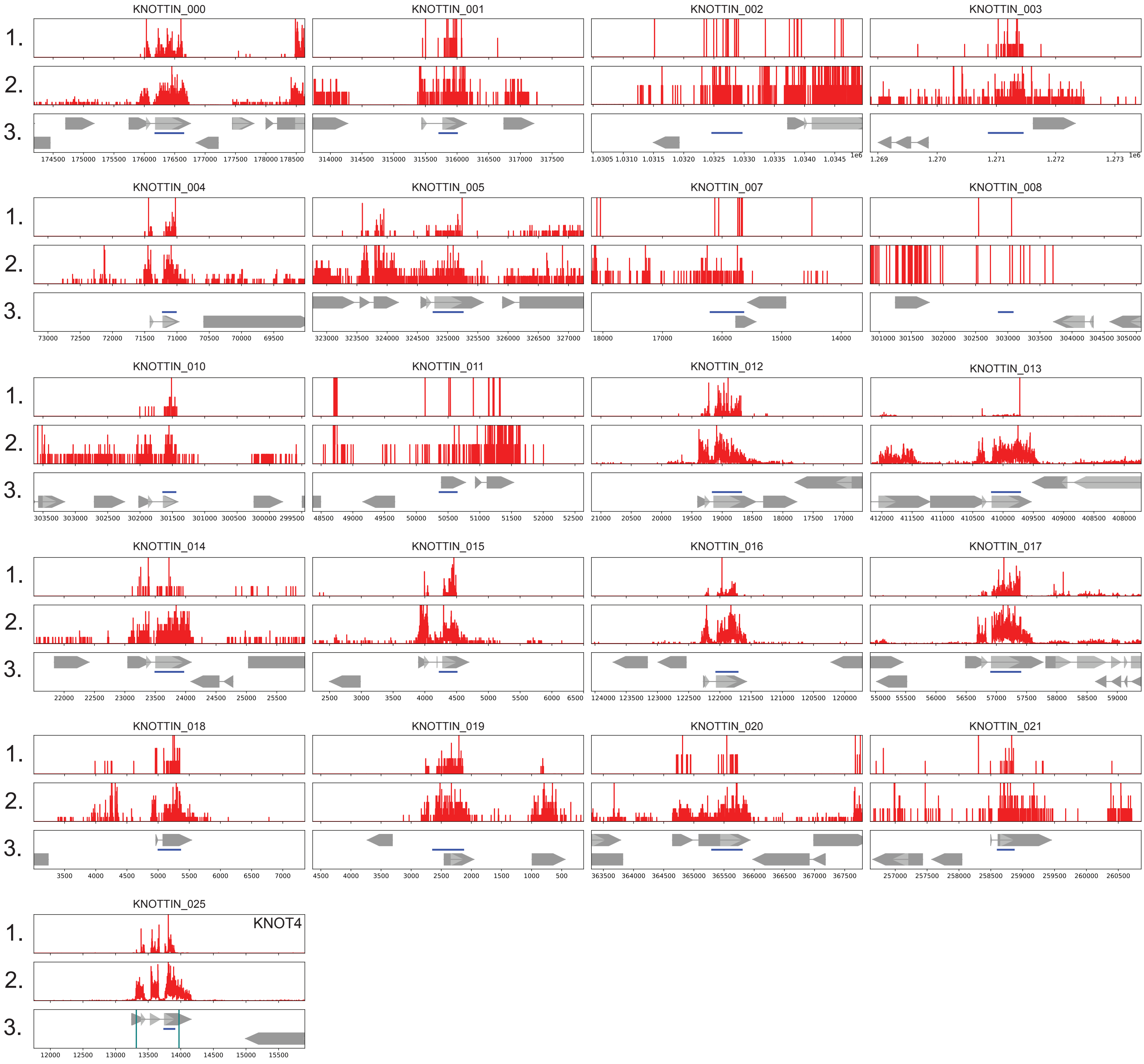


Figure S3. Ribosome profiling and transcriptomics results for the remaining 21 putative knottins.

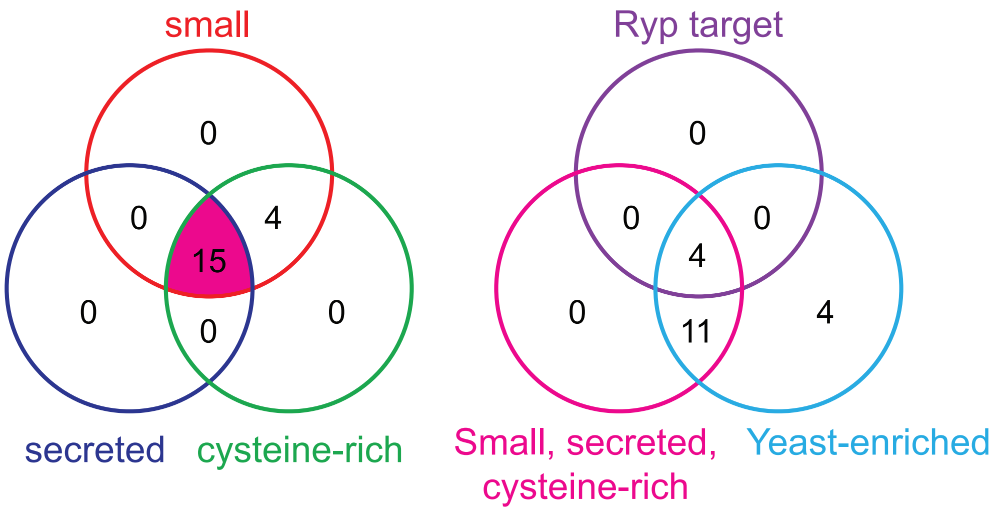


Figure S4. Effector criteria applied to the 25 putative knottins.

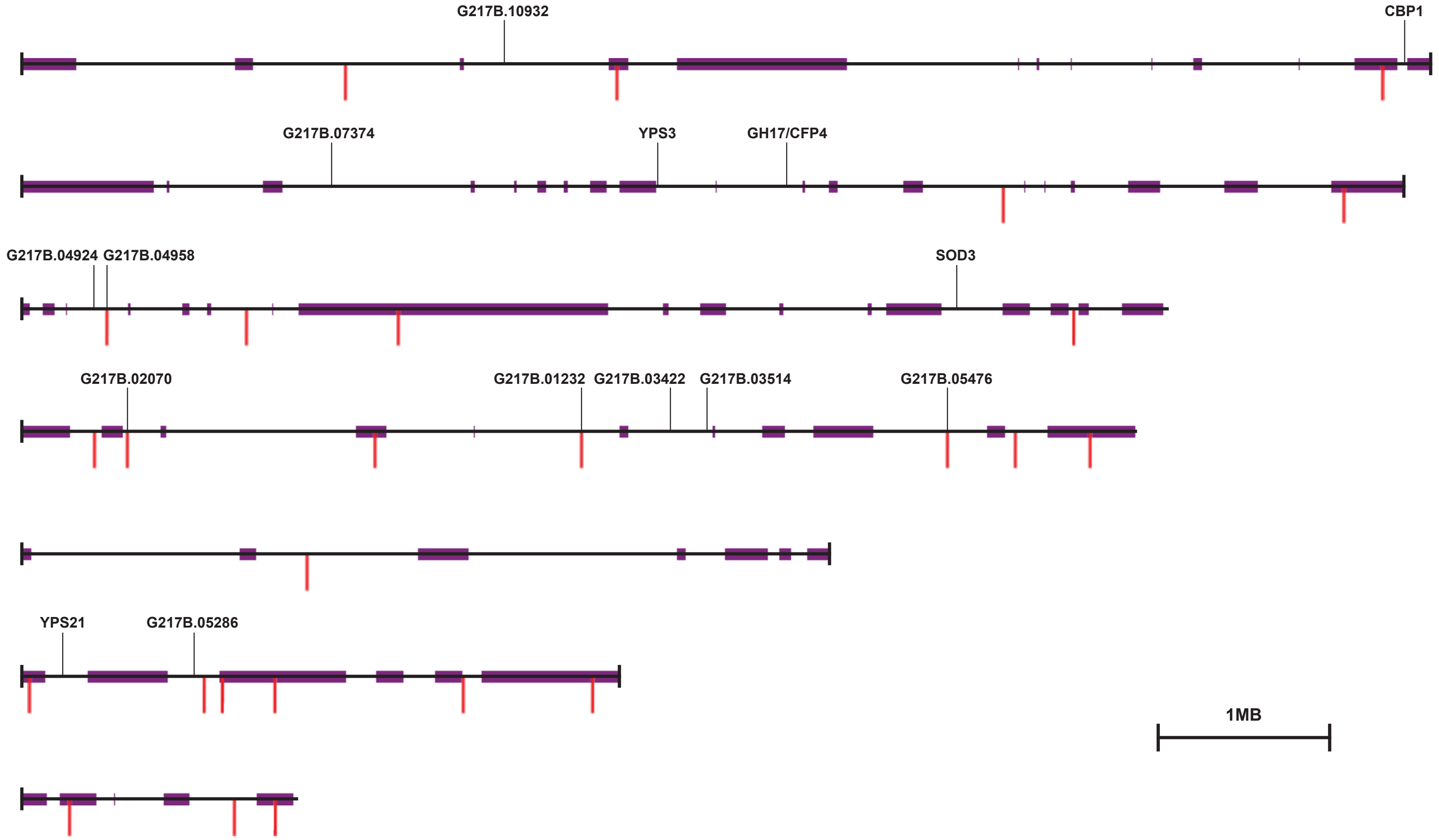


Figure S5. Nanopore sequencing reveals random distribution of knottins and putative virulence factors throughout the G217B *Histoplasma* genome.

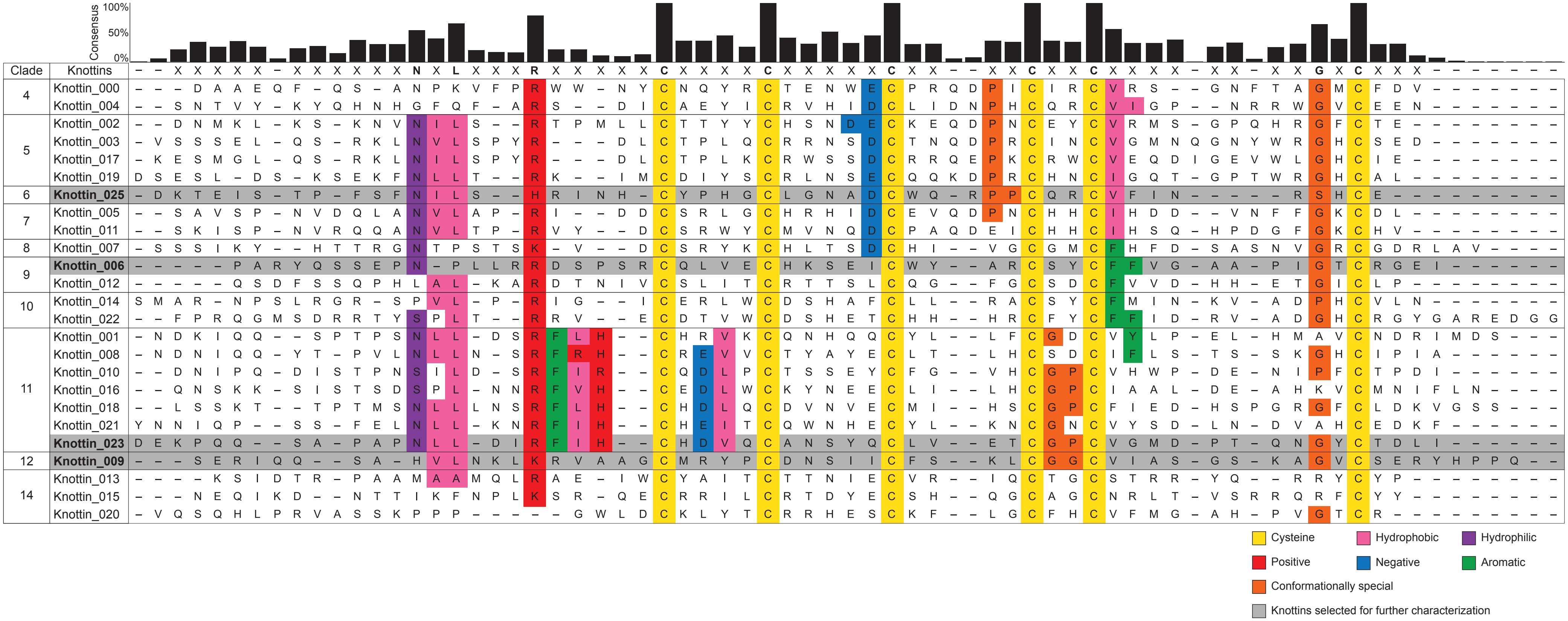


Figure S6. Protein alignment of the 25 putative knottins.

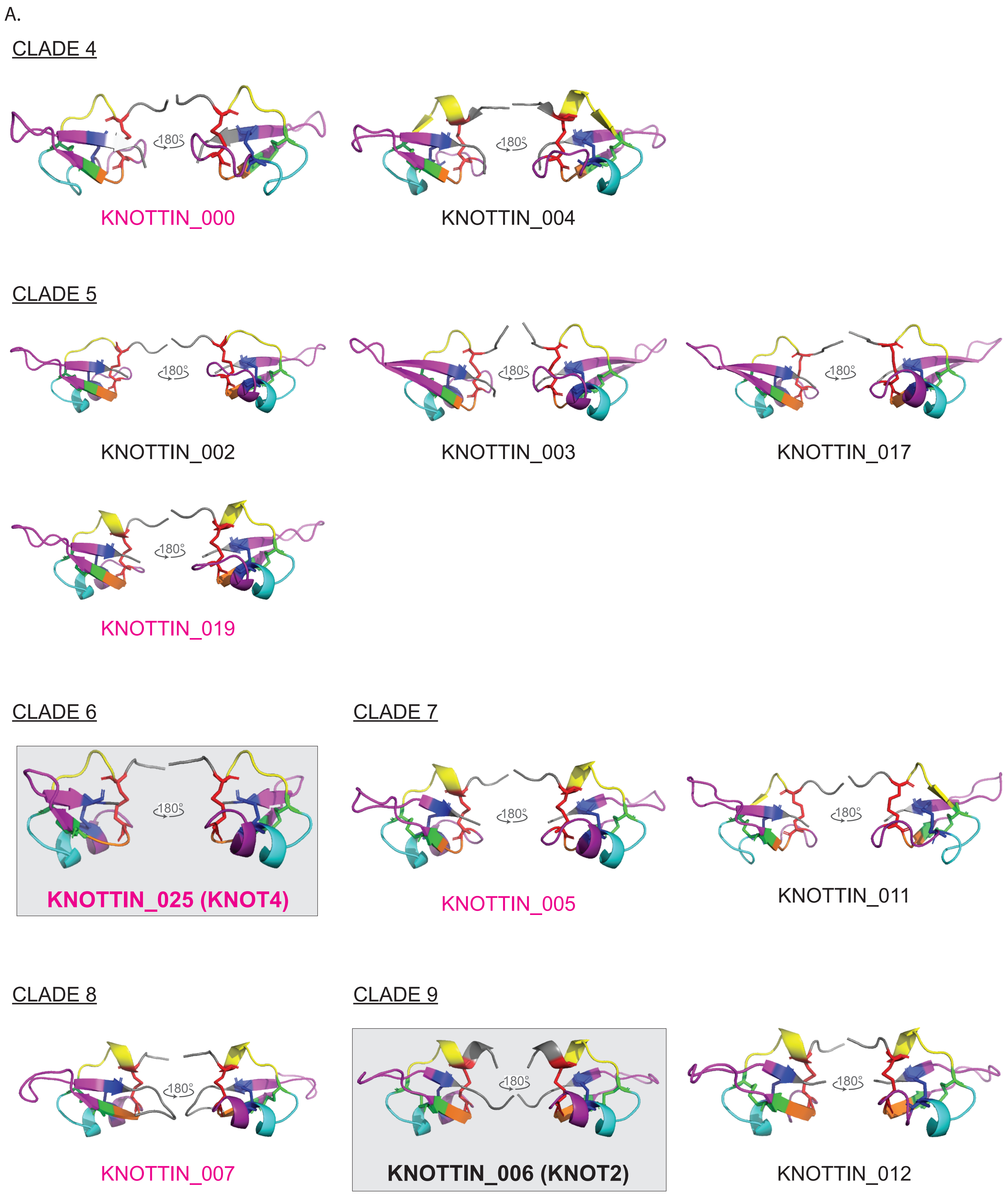
A.
CLADE 4
180°
KNOTTIN_000
180°
KNOTTIN_004
CLADE 5
180°
KNOTTIN_002
180°
KNOTTIN_003
180°
KNOTTIN_017
180°
KNOTTIN_019
CLADE 6
180°
KNOTTIN_025 (KNOT4)
CLADE 7
180°
KNOTTIN_005
180°
KNOTTIN_011
CLADE 8
180°
KNOTTIN_007
CLADE 9
180°
KNOTTIN_006 (KNOT2)
180°
KNOTTIN_012

A. continued

CLADE 10

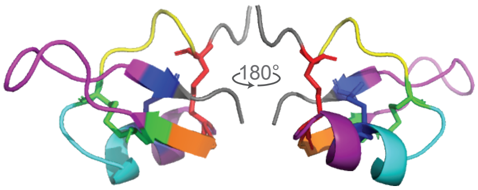


KNOTTIN_014

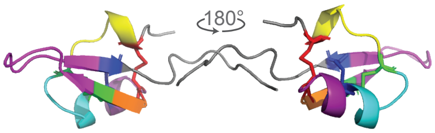


KNOTTIN_022

CLADE 11

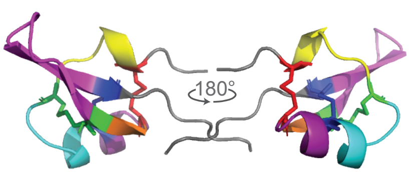


KNOTTIN_001

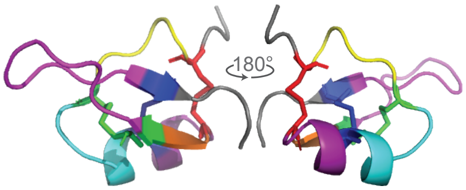


KNOTTIN_008

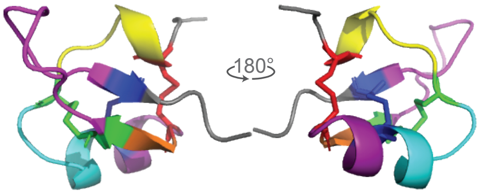


KNOTTIN_010

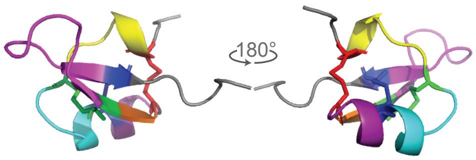


KNOTTIN_016

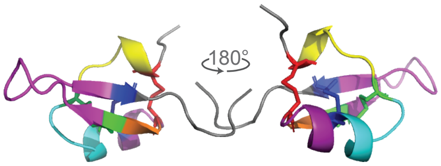


KNOTTIN_018

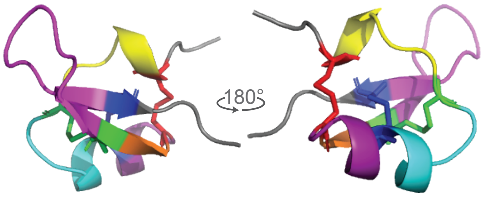


KNOTTIN_021

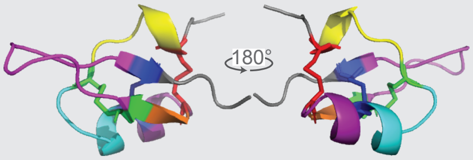


**KNOTTIN_023 (KNOT3)**

CLADE 12

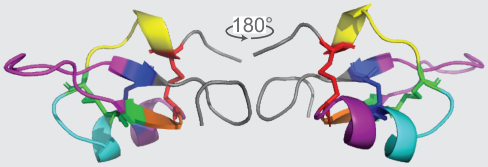


**KNOTTIN_009 (KNOT1)**

CLADE 14

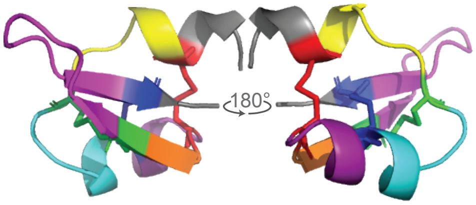


KNOTTIN_013

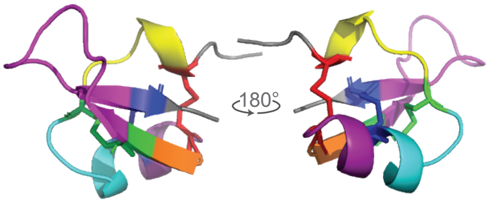


KNOTTIN_015

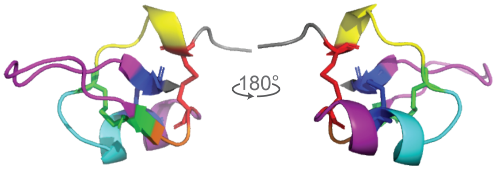


KNOTTIN_020

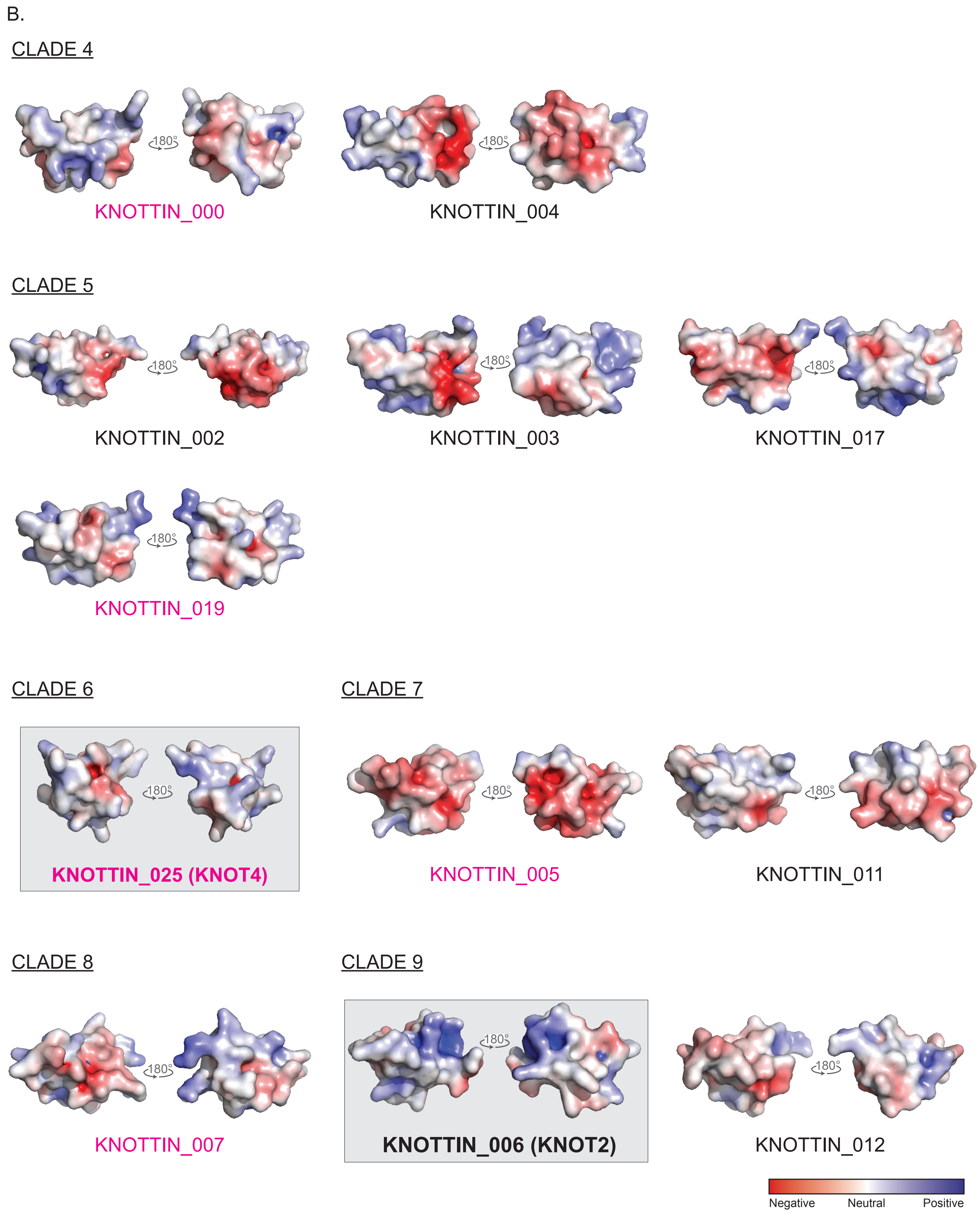
B.
CLADE 4
180°
KNOTTIN_000
KNOTTIN_004
CLADE 5
KNOTTIN_002
KNOTTIN_003
KNOTTIN_017
KNOTTIN_019
CLADE 6
KNOTTIN_025 (KNOT4)
CLADE 7
KNOTTIN_005
KNOTTIN_011
CLADE 8
KNOTTIN_007
CLADE 9
KNOTTIN_006 (KNOT2)
KNOTTIN_012
Negative
Neutral
Positive

B. continued

CLADE 10

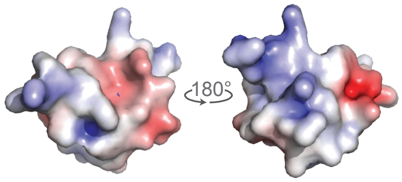


KNOTTIN_014

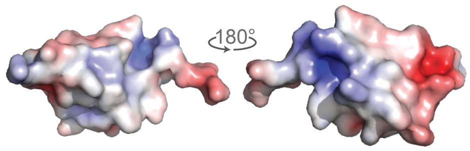


KNOTTIN_022

CLADE 11

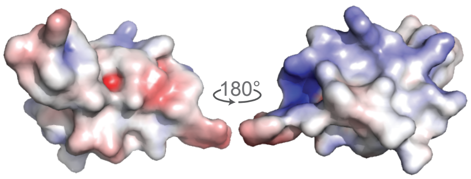


KNOTTIN_001

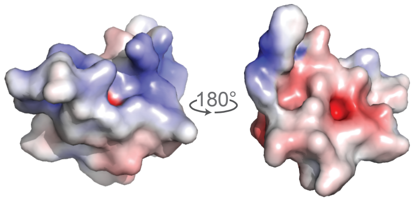


KNOTTIN_008

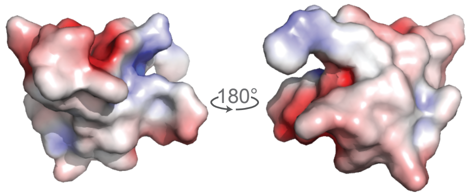


KNOTTIN_010

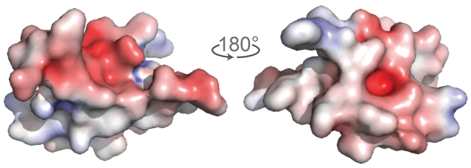


KNOTTIN_016

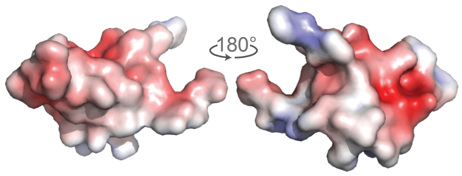


KNOTTIN_018

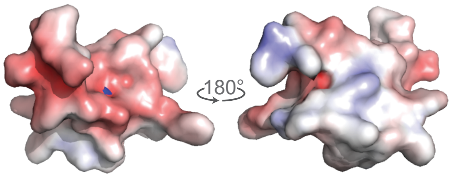


KNOTTIN_021

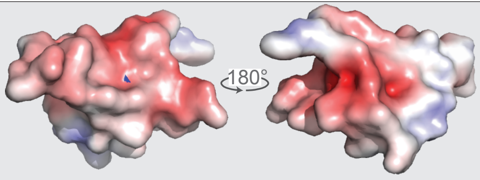


**KNOTTIN_023 (KNOT3)**

CLADE 12

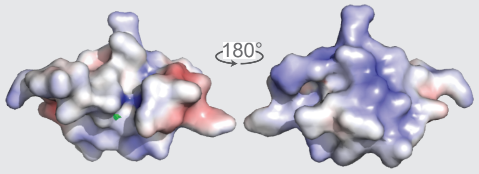


**KNOTTIN_009 (KNOT1)**

CLADE 14

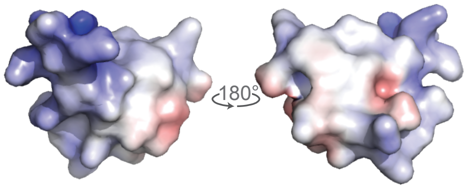


KNOTTIN_013

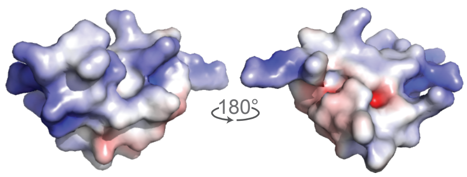


KNOTTIN_015

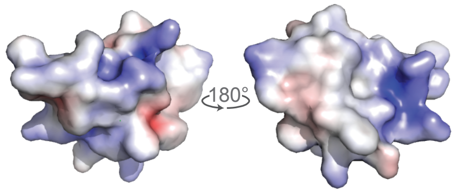


KNOTTIN_020

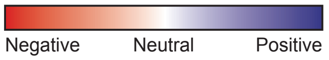


Figure S7. Predicted protein structures of all putative knottin.

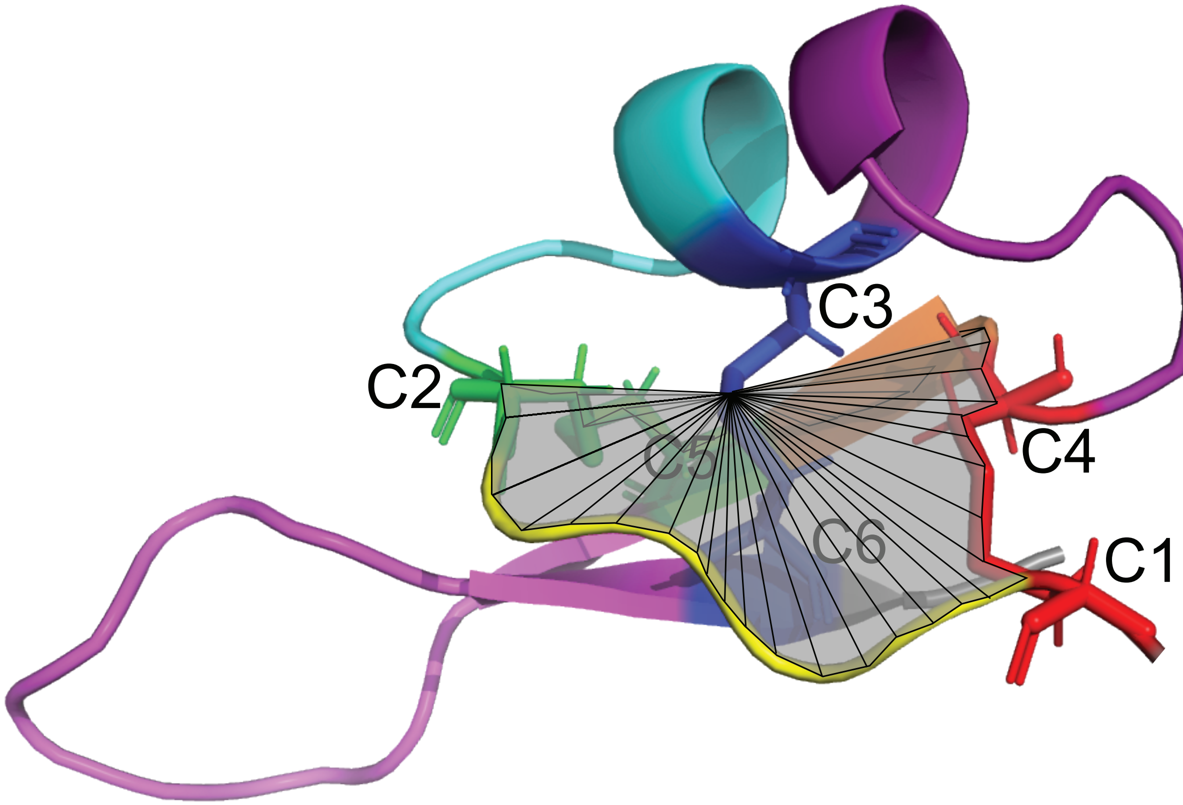


Figure S8. Visual representation of the KNOTTIN_TOPOLOGY_SCAN algorithm.

| Clade | Species and knottin | Protein Sequence |
|---|---|---|
| 6 | **G217B_025** | FSFNILSHRINHCYPHGCLGNADCWQRPPCQRCVFINRSHCE ← Knot4 |
| | G186AR_005 | FSFNILSHRMNHCYPFGCLGNADCWRRPPCQRCVFANRSHCE |
| | H88_024 | FSFNILSHRMDHCYPFGCLGHADCWRRPPCQHCVFANRSHCE |
| | H143_015 | FSFNILSHRMDHCYPFGCLGHADCWRRPPCQHCVFANRSHCE |
| | WU24_014 | ISFNILSHRMDHCFPYGCLGHADCWRRPPCQRCVFVNRSHCE |
| | Tmu_007 | FSFDILSHRMNHCYPFGCLGHADCWRRPPCQRCVFANRSHCE |
| 9 | **G217B_006** | N-PLLRRDSPSRCQLVECHKSEICWYARCSYCFFVGAAPIGTCRGEI ← Knot2 |
| | G186AR_004 | N-PLLQREAPGRCQEVGCHKSELCWYSRCTYCLFVGASILGTCRGER |
| | H88_011 | N-PLLQRDAPSRCQHVECHKSEICWYARCSYCFFVGASPIGTCRGER |
| | H143_006 | N-PLLQRDAPSRCQHVECHKSEICWYARCSYCFFVGASPIGTCRGER |
| | WU24_003 | N-PLLRRDSPSHCLHVECHKSEICWYSHCSWCFFVGASPIGTCRGER |
| | Tmu_022 | N-PLLQRNAPGRCQHVECHKSEICWYSRCSYCFFVGAAPIGSCRGEI |
| | **G217B_012** | HLALKARDTNIVCSLITCRTTSLCQGFGCSDCFVVDHHETGICL--P |
| | G186AR_020 | HLALKARDTNVVCSMIDCRTTSLCRGFGCSECFVVDHHKTGICL--P |
| | H88_025 | HLALKARDTNIVCSLITCRTTSLCQGFGCSDCFVVDHHETGICL--P |
| | H143_002 | HLALKARDTNIVCSLITCRTTSLCQGFGCSDCFVVDHHETGICL--P |
| | Tmu_019 | HLALKARDMNVVCSLITCRTTSLCQGFGCSECFVVDHHETGICL--P |
| 11 | Tmu_002 | MSNLLLNSRFLRCHDLLCDANIECLIHSCGPCFFPEDSPGKGYCLDKVDS------S |
|  | G186AR_012 | MSNLLLNSRFLRCHDLPCDVNLECLVHSCGPCFFPEDSPGKGYCLDKVDS------S |
|  | H88_019 | MSNLLLNSRFLHCHDIPCDANIECMVHSCGPCYFPEDSPGKGYCLDKVGS------S |
|  | H143_017 | MSNLLLNSRFLHCHDIPCDANIECMVHSCGPCYFPEDSPGKGYCLDKVGS------S |
|  | **G217B_018** | MSNLLLNSRFLHCHDLQCDVNVECMIHSCGPCFIEDHSPGRGFCLDKVGS------S |
|  | **G217B_023** | APN-LLDIRFIHCHDVQCANSYQCLVETCGPCVGMD-PTQNGYCTD---L------I ← Knot3 |
|  | G186AR_000 | APN-LLDIRFIHCHDVECANSYQCLIQACGPCVGMD-PTQNGVCTD---L------L |
|  | H88_000 | APN-LLDIRFIHCHDVQCANSYQCLMQLCGPCVSMD-PTQNGVCTD----------L |
|  | H143_008 | APN-LLDIRFIHCHDVQCANSYQCLMQLCGPCVSMD-PTQNGVCTD----------L |
|  | WU24_020 | ASN-LLDIRFIHCHEVECANSYQCLIQSCGPCVGMD-PIKNGVCTD---L------H |
| | Tmu_014 | APN-LLDIRFIHCHDVQCANSYQCLIVSCGPCVGMD-PTKDGVCTD---LLDDQLMG |
|  | **G217B_016** | TSDSPLNNRFVHCEDLWCKYNEECLILHCGPCIAAL-DEAHKVCMNI-FL------N |
|  | G186AR_014 | TSNLALNNRFVHCEDLWCEYNEECLVIHCGPCVATL-EEDEKVCMNI-IM-----K |
|  | H143_005 | TSNLALNNRFVHCEDLWCEYNEECLVLQCGPCIATL-EEDEKVCMNI-FL------N |
|  | H88_001 | TSNLALNNRFVHCEDLWCEYNEECLVLQCGPCIATL-EEDEKVCMNI-FL------N |
|  | Tmu_008 | TSNLALNNRFVHCDDLWCEYNEECLVLNCGPCIASL-NEDQKVCMNI-FL------K |
|  | WU24_019 | TPNSILDSRFILCQDLPCTSSEYCFNFHCGPCVHWS-DEDIPLCTP---D------F |
|  | **G217B_010** | TPNSILDSRFIRCQDLPCTSSEYCFGVHCGPCVHWP-DENIPFCTP---D------I |
|  | Tmu_018 | TSNLILDSRFIWCQDLHCTSSEYCFNAHCGPCVHWP-DETIPFCTP---D------L |
|  | G186AR_002 | TPNSILDSRFIRCQDLPCTSSGYCFNSHCGPCVHWP-DENIPFCTP---D------H |
|  | H143_004 | PSN-LLESRLVHCHDISCQYHQQCYLLYCGDCIFLP-SRNYAICNDRIVH------R |
|  | H88_003 | PSN-LLESRLVHCHDISCQYHQQCYLLYCGDCIFLP-SRNYAICNDRIMH------R |
|  | G186AR_013 | PSN-LLESRLLRCHRISCQNHQQCYLLYCGDCVFLP-SLNFAVCNDRIVH------R |
|  | **G217B_001** | PSN-LLDSRFLHCHRVKCQNHQQCYLLFCGDCVYLP-ELHMAVCNDRIMD------S |
|  | WU24_005 | PSN-LLESRLLYCHRVHCRNHQQCYLLYCGDCTFLP-SLNMAVCDDRIMN------P |
|  | Tmu_016 | PSN-LLESRLLRCHEISCQYSQHCYLLYCGDCIVLP-Q-QVARCNDIIVH------R |
|  | **G217B_021** | ELN-LLKNRFIHCHEITCQWNHECYLKNCGNCVYSD-LNDVAHCEDK---------F |
|  | H143_021 | ILN-LLKNRFIHCHEVTCKWHHECYLQKCGNCVYSD-LNDVAHCEDK---------I |
|  | H88_006 | ILN-LLKNRFIHCHEVTCKWHHECYLQKCGNCVYSD-LNDVAHCEDK---------I |
|  | G186AR_021 | ILN-LLKNRFIHCHEVTCKWHHECYLRKCGNCVYSD-LNDVAHCEDK---------F |
|  | Tmu_011 | VLS-LLKNRFIHCHEITCKWHHECYLQKCGNCVYSD-LNDMAHCEDK---------F |
|  | Tmu_015 | ----VLNSRFRYCRDIVCTYTYECLTLHCSDCILLS-TSSKGHCIP---I------A |
|  | WU24_006 | VLN-LLNSRFRYCREIVCTYAYECLTLHCSDCIFLS-TSSKGHCIP---I------D |
|  | G186AR_022 | VLN-LLNSRFRHCREVVCTYAYECLILHCSDCIFLS-TSSKGHCIP---I------A |
|  | H88_013 | VLN-LLNSRFRHCREVVCTYAYECLTLHCSDCIFLS-TSSKGHCIP---I------A |
|  | **G217B_008** | VLN-LLNSRFRHCREVVCTYAYECLTLHCSDCIFLS-TSSKGHCIP---I------A |
|  | H143_014 | VLN-LLNSRFRHCREVVCTYAYECLTLHCSDCIFLS-TSSKGHCIP---I------A |
| 12 | **G217B_009** | HVLNKLKRVAAGCMRYPCDNSIICFSKLCGGCVIASGSKAGVCSERYHPPQ ← Knot1 |
| | G186AR_001 | QVLNKLKRVAAGCMRYPCDNSIICFSKLCGGCVIASGSKAGVCSERYHPPR |
| | H88_010 | QVLNKLKRAAAGCMRYPCDNSIICFSKLCGGCVIASGSKAGVCSERYHPPR |
| | H143_011 | QVLNKLKRAAAGCMRYPCDNSIICFSKLCGSCVIASGSKAGVCSERYHPPR |
| | WU24_001 | HVLNKLKRGPSGCMRYPCDNSIICYSKLCGSCVFPSGSKAGVCSPRYHPPR |
| | Tmu_009 | HVLNKLKRVAAGCMRYPCDNSIICFSKFCGGCVIASGSKAGVCSERYHPPR |

Cysteine | Hydrophobic | Hydrophilic
Positive | Negative | Aromatic
Conformationally special

Figure S9. Knot1-4 protein alignment across different *Histoplasma* species.

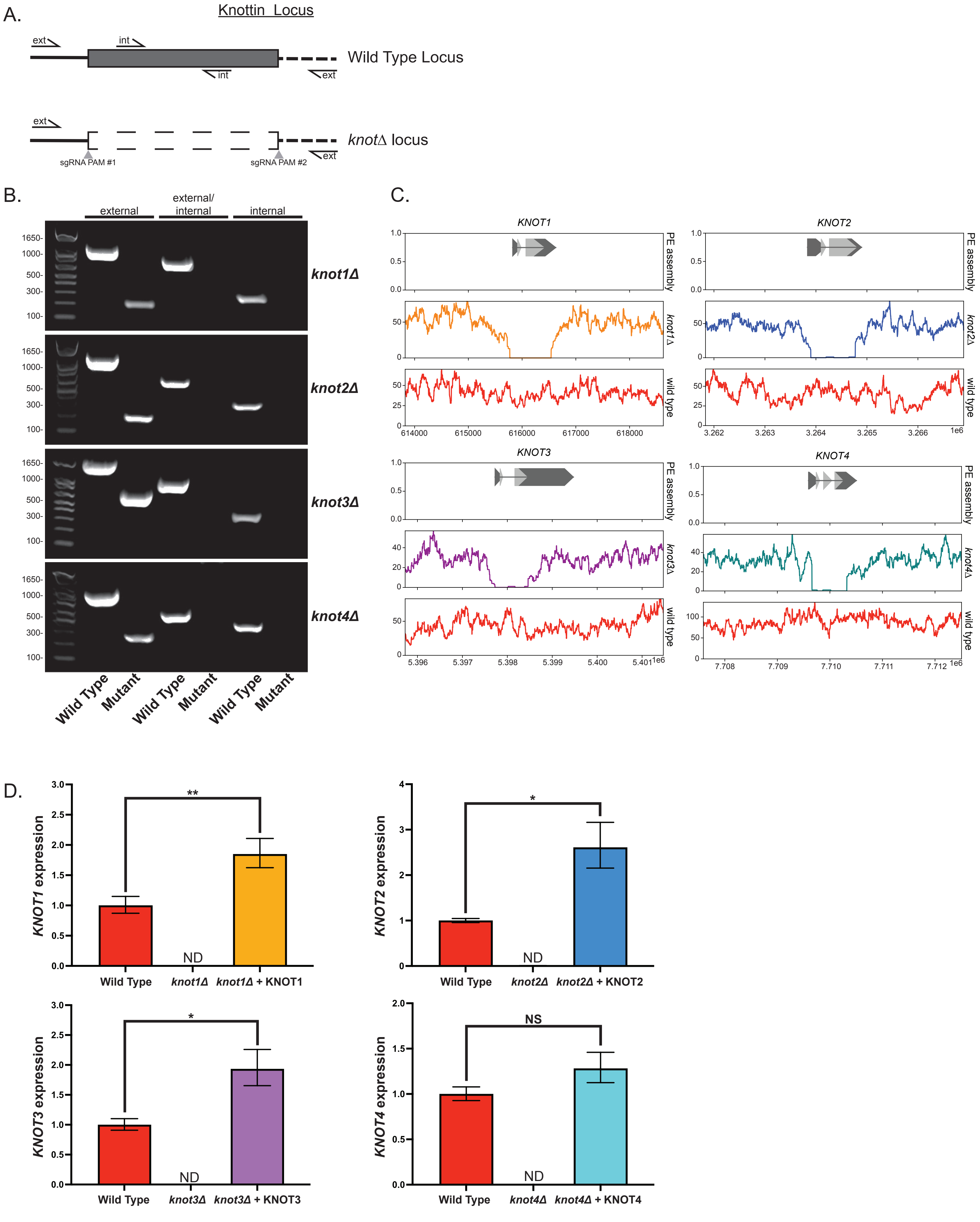


Figure S10. Deletion and complementation of *KNOT1-4* and validation of strains.

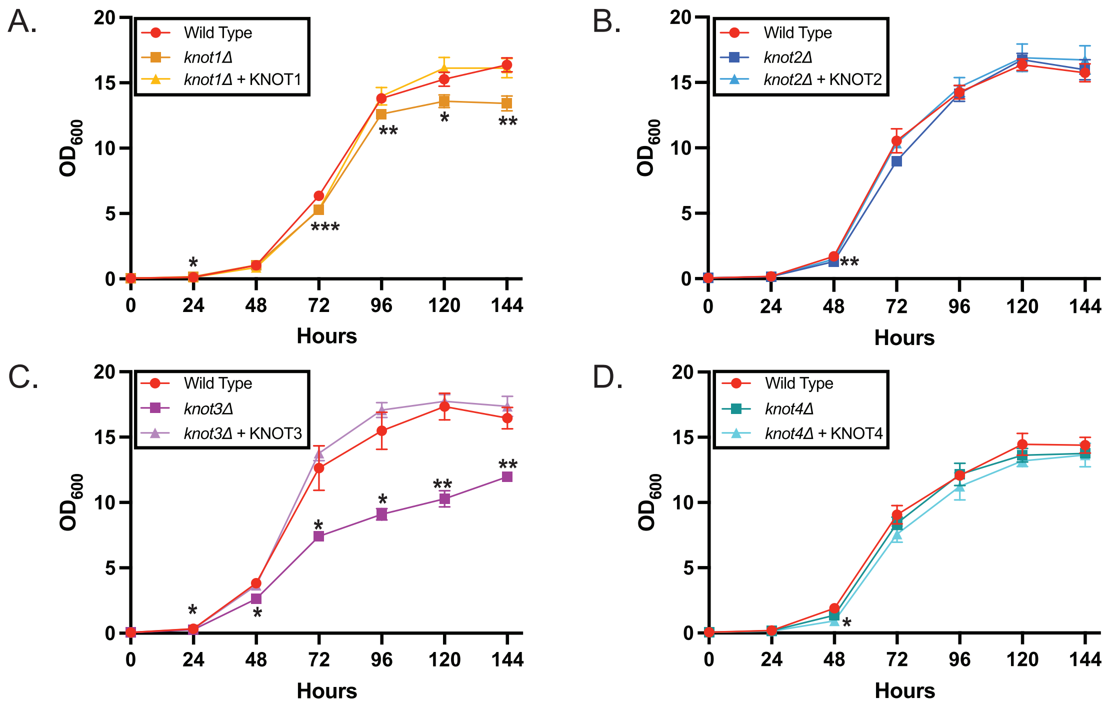


Figure S11. *In vitro* growth curves reveal Knot3 is required for optimal growth.

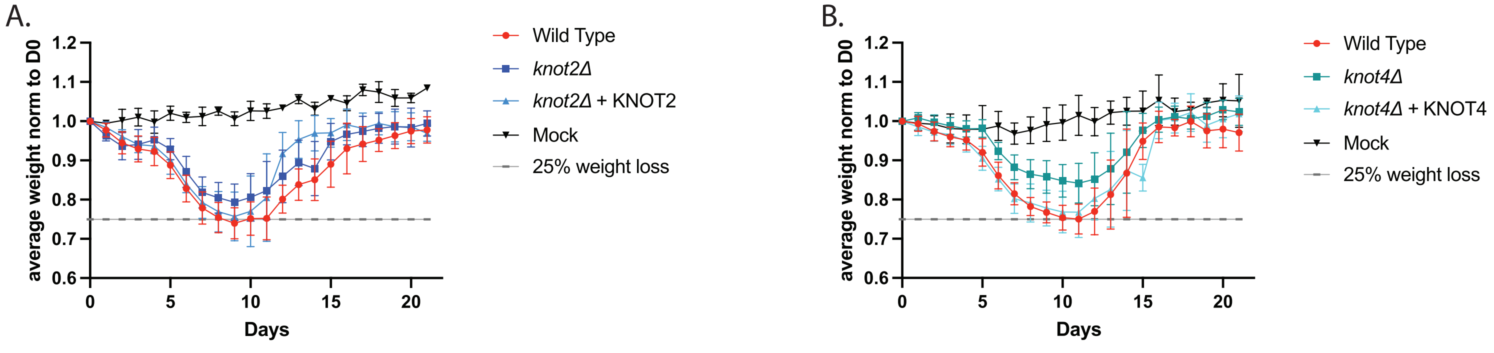


Figure S12. Mice infected with *knot2*Δ or *knot4*Δ mutant lose weight but do not succumb to infection.
